# Linker histone H1 functions as a liquid-like glue to organize chromatin in living human cells

**DOI:** 10.1101/2025.03.05.641622

**Authors:** Masa A. Shimazoe, Jan Huertas, Charles Phillips, Satoru Ide, Sachiko Tamura, Stephen Farr, S. S. Ashwin, Masaki Sasai, Rosana Collepardo-Guevara, Kazuhiro Maeshima

## Abstract

Linker histone H1, the most abundant chromatin protein, condenses chromatin, modulates DNA transactions such as transcription and DNA replication/repair, and participates in differentiation, development and tumorigenesis. While recent studies indicate that nucleosomes are clustered as condensed chromatin domains in higher eukaryotic cells, how histone H1 mechanically condenses chromatin remains unclear. Here, using a combination of direct visualization of single-H1 molecules in living human cells and multiscale molecular dynamics simulations, we demonstrate that the majority of H1 behaves like a liquid inside chromatin domains, rather than binding stably to nucleosomes as suggested by the traditional model. H1 functions as a liquid-like “glue”, mediating dynamic multivalent electrostatic interactions between nucleosomes within chromatin domains. Consistently, rapid-depletion of H1.2 leads to decondensed chromatin domains both in cells and *in silico*. Our findings suggest that the H1 “glue” condenses chromatin domains while keeping them fluid and accessible, thereby supporting essential DNA transactions.

## Introduction

How is the long string of nucleosomes—where DNA is wrapped around core histones (2 copies of H3, H4, H2A and H2B) (*1–4*) —organized into chromatin in living cells (*5, 6*)? A growing body of evidence indicates that chromatin is a highly dynamical and variable structure, and is folded irregularly into condensed chromatin domains in higher eukaryotic cells (*7–17*). Genome-wide genomics analyses, such as Hi-C (*18*), have also revealed the presence of chromatin domains with distinct epigenetic marks (*19–22*).

What factors contribute to the condensation of chromatin into domains? A key contributor is nucleosome–nucleosome interactions mediated by histone tails (*8, 12, 23–27*), facilitated by cations such as Mg²⁺ (*28, 29*). Another critical factor is the linker histone H1, which condenses chromatin both *in vitro* and *in vivo* (*30–34*) and promotes chromatin phase separation *in vitro* (*27*). H1 is the most abundant chromatin-binding protein (*35, 36*). It is highly conserved across eukaryotes and forms a large family with 11 distinct subtypes in both humans and mice (*37*).

Structurally, H1 is a tripartite protein consisting of a well-defined and conserved globular domain (GD) with approximately 80 amino acid residues, flanked by a short, unstructured N-terminal domain (NTD) containing 20–35 residues, and a long, intrinsically disordered C-terminal domain (CTD) with around 100 residues (Fig. S1A). The CTD comprises about half of the H1 sequence across all subtypes and is highly positively charged, with lysine residues making up approximately 40% of its amino acid composition. It has a high affinity for nucleosomal and linker DNA and is essential for chromatin condensation (*31–34*).

H1 binding to the nucleosome induces structural changes in chromatin, thus regulating transcription and other DNA-dependent processes (*36, 38*). Consistently, early mouse knockout studies revealed that embryos with reduced H1 levels cease development at mid-gestation (*39*). H1 functions in gene regulation through 3D genome organization (*40*), and depletion of H1 leads to perturbations of gene expression, particularly in constitutive heterochromatin in mouse embryonic stem cells (*41*). Furthermore, mutations in genes encoding several H1 isoforms are highly recurrent in B-cell lymphomas (*42*). These findings suggest pivotal roles for H1 in development, differentiation, and tumorigenesis through its regulation of higher-order chromatin organization.

How does linker histone H1 condense chromatin into domains within the cell? The classical textbook model has posited that H1 stably binds to the nucleosome dyad (the central base pair of nucleosomal DNA) and stabilizes the 30-nm chromatin fiber (*43–45*). Binding modes of H1 around the nucleosome dyad have been well characterized (*45–50*). However, recent evidence indicates that the 30-nm fiber is not the fundamental chromatin structure in living cells (*25, 51–53*). Instead, chromatin is organized into irregular, dynamic chromatin domains (*54*).

Fluorescence Recovery After Photobleaching (FRAP) studies have shown that H1 binds to chromatin in a highly dynamic, rather than stable, manner (*55–59*). Although the majority of H1 molecules are bound to chromatin at any given moment, this binding is transient, with residence times of approximately one minute. These findings underscore the need for a new molecular mechanism to explain how H1 interacts with nucleosomes and regulates chromatin structure. Additionally, chromatin is highly charged and its structure can vary greatly depending on the surrounding environment, such as the presence of cations, macromolecular crowding (*60, 61*), or fixation with chemical crosslinking (*62*). There is also a need for new technology to investigate linker histone H1 behavior in chromatin in living cells.

To address these challenges, we employed an interdisciplinary approach combining single-molecule imaging and tracking techniques (*63–67*), multiscale molecular dynamics (MD) simulations of chromatin (*68–70*), and rapid protein depletion technology (*71*). Single-molecule imaging and tracking enable the investigation of molecular behavior in living cells, while multiscale simulations provide mechanistic insights into chromatin organization across different resolutions.

By integrating these approaches, we demonstrate that H1 acts as a liquid-like “glue” for chromatin. In this role, H1 moves dynamically around and between nucleosomes within chromatin, effectively screening electrostatic repulsion between multiple DNA linkers. This dynamic binding enables H1 to condense chromatin while simultaneously fostering DNA fluctuations and a diverse range of inter-nucleosome distances and orientations, thereby promoting an irregular liquid-like chromatin organization.

## Results

### Computational modeling suggests that linker histone H1 acts as a liquid-like “glue” for chromatin

To gain insights into the behavior of linker histone H1 in chromatin, we first conducted unbiased coarse-grained molecular dynamics (MD) simulations of single nucleosomes and 12-mer nucleosome arrays to examine the binding of H1 in both contexts (Movies 1-2)(Fig.1A, left and center). We used our chemically-specific coarse-grained model (*72*), in which each amino acid of the histones was represented by one bead and each DNA base pair by one ellipsoid and one point charge per phosphate (see Methods). We focused on H1.2, one of the most abundant linker histone variants. Importantly, in these simulations, the H1.2 protein was free to dynamically bind and unbind from the nucleosome, rather than being permanently fixed to the dyad. This dynamical binding/unbinding behavior is regulated by two factors: the interplay of electrostatic and hydrophobic interactions between amino acids and DNA phosphates, and the large ensemble of configurations sampled by H1.2.

**Fig. 1.**
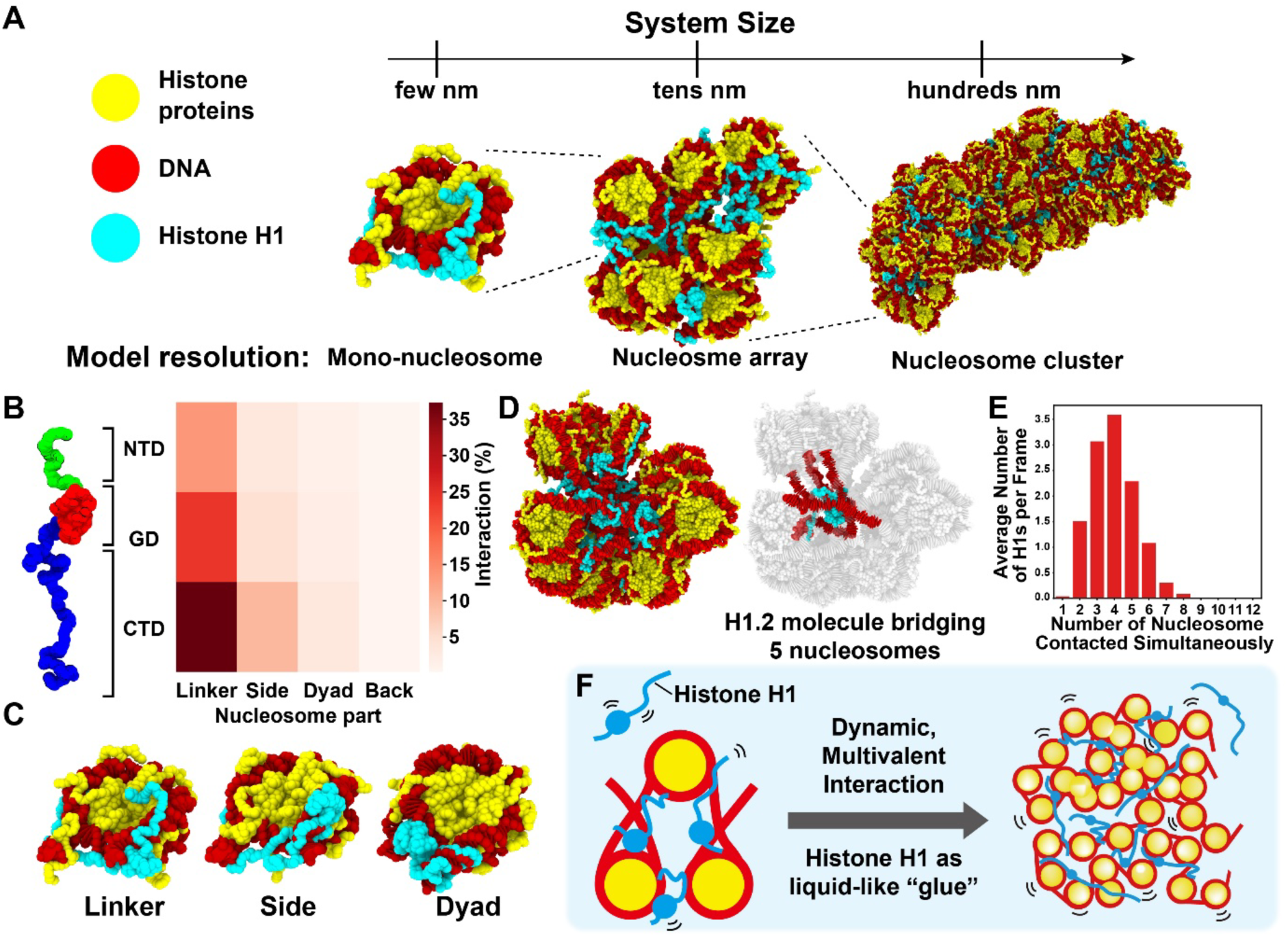
Multiscale computational modeling and a new binding model of linker histone H1. **(A)** Scheme for multiscale computational modeling. A single nucleosome, a 12-mer nucleosome array with a nucleosome repeat length (NRL) of 195-bp, and a 108-mer nucleosome cluster (195 bp NRL) are shown. All simulations retain amino acid-level resolution. **(B)** Region-specific contact map between H1.2 and nucleosomes for a 12-mer nucleosome array (195 bp NRL). H1.2 (y-axis): NTD, GD, CTD regions; Nucleosome (x-axis): Linker, Side, Dyad, and Back regions. **(C)** Examples of interactions between H1.2 and a nucleosome. **(D)** A representative configuration of a 12-mer nucleosome array (195 bp NRL) with H1.2 (left). One H1.2 can bridge multiple nucleosomes at a time (right) (see also Movies S1-S2). **(E)** Histogram of the number of nucleosomes contacted simultaneously by a single H1.2. **(F)** A new model: linker histone H1 dynamically interacts with nucleosomes through multivalent interactions (left) and has a liquid-like “glue” activity to condense chromatin domains (right).

From these simulations, we calculated the relative frequencies of contacts between the different structural regions of H1.2 (Fig. 1B, left) — its N-terminal domain (NTD), the globular domain (GD), and its C-terminal domain (CTD) — and the DNA of the nucleosome. The DNA was categorized into four regions: the side, back and dyad of the nucleosomal DNA, and the linker DNA (Fig.1C). Interestingly, although the dyad has long been considered the primary H1 binding site for the GD within the nucleosome, our contact map shows otherwise. In the mono-nucleosome, all H1.2 domains bind preferentially to the nucleosome side (Fig. S1B), while in the 12-mer nucleosome array all domains more frequently interact with the linker DNA, followed by the nucleosome side (Fig. 1B, right), showing no strong preference for the dyad in either case.

Because unbiased MD simulations can fail to sufficiently explore the conformational landscape of large and heterogeneous systems like chromatin, we next performed Temperature Replica Exchange Molecular Dynamics (T-REMD) simulations of 12-mer nucleosome arrays, with one H1.2 per nucleosome, to enhance sampling of H1.2 configurations and assess the impact of H1.2 on inter-nucleosomal interactions (Fig. S1C). Analysis of the frequency of interactions between different H1.2 molecules and the various nucleosomes in the array (Fig. 1D) indicated that each H1.2 molecule forms weak and transient contacts with multiple nucleosomes simultaneously (Figs.1D-E), continuously exchanging partners (Figs. S1D-F). Most commonly, a single H1.2 molecule contacts the DNA of two distinct nucleosome cores (Fig. S1E), and interacts with the linker and nucleosomal DNA across four different nucleosomes (Fig. 1E). Remarkably, in some cases, an H1.2 molecule embedded deep within the nucleosome array bridges as many as 8 separate nucleosomes (considering linker and nucleosomal DNA; Fig. 1E). This exceptional capacity of H1 to form multivalent interactions with nucleosomes drives the emergence of a dynamic, irregular, and compact cluster of nucleosomes (Fig. S1G), which behaves like a “fluid”.

Based on these findings, we propose a “liquid-like glue” model for linker histone H1 function (Fig. 1F). In this model, each H1 molecule simultaneously engages several nucleosomes through its GD and intrinsically disordered CTD. These interactions are highly transient: the GD and CTD frequently disengage from one nucleosome and rebind to another in an uncoordinated, stochastic, manner (Fig. S1F). This lack of coordination is critical: while one domain may transiently unbind, the other typically remains engaged, thereby maintaining the overall association of H1 with the chromatin. Fully coordinated unbinding of all domains is rare—but statistically plausible—and would lead to complete dissociation of H1 from the chromatin. By contacting multiple nucleosomes simultaneously via its different domains and frequently exchanging nucleosome binding partners, H1 facilitates condensation of chromatin while ensuring its fluidity.

### Single-H1 imaging in live human cells

To investigate H1 behavior in chromatin and validate our model, we performed single-H1 molecule imaging in live human cells. We used the CRISPR/Cas9 system (*73*) to introduce a HaloTag sequence at the C-terminus of the endogenous histone H1.2 gene in human Retinal Pigment Epithelial (RPE-1) cells (Fig. 2A) (*74*). The HaloTag allowed for visualization with tetramethylrhodamine (TMR; Fig. 2B) or other Janelia Fluor (JF) dyes (*75*). Cell clones stably expressing H1.2-Halo were labeled with TMR and isolated using fluorescence-activated cell sorting (FACS; Fig. S2A)(*76*). Proper mono-allelic HaloTag insertion and expression of the H1.2-Halo were confirmed by polymerase chain reaction (PCR) and western blotting (Figs. 2C and S2B-C). The distribution of H1.2-Halo in RPE-1 cells resembled the pattern observed with 4,6-diamidino-2-phenylindole (DAPI) staining (Pearson’s correlation = 0.88, Figs. 2B and S2D), indicating genome-wide localization including heterochromatin. Stepwise-salt washing of nuclei isolated from H1.2-Halo expressing cells demonstrated that the biochemical properties of H1.2-Halo closely matched those of native H1.2 (Fig. S2E), confirming appropriate chromatin association. Additionally, computational modeling of chromatin incorporating H1.2-Halo produced a region-specific contact map and sedimentation coefficient (S) comparable to those obtained without the HaloTag (Figs. S1G-5 and S2F), further validating our tagging approach.

**Fig. 2.**
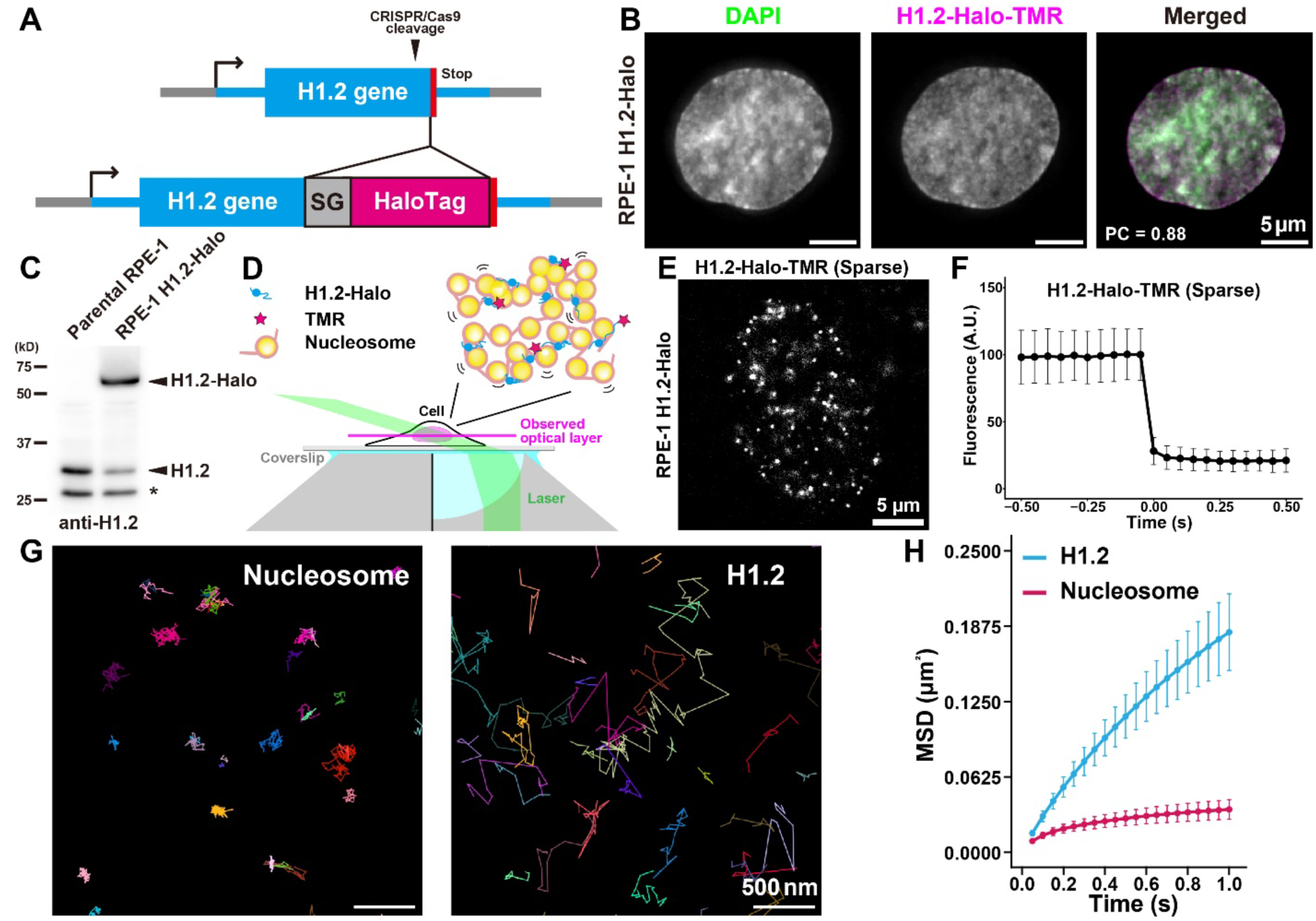
RPE-1 cells stably expressing H1.2-HaloTag and single-molecule H1.2 imaging. **(A)** Schematic of CRISPR/Cas9-mediated genome editing for inserting the HaloTag at the C-terminus of the H1.2 gene. SG: linker (GGGGS x3); Stop: stop codon. **(B)** Representative images of RPE-1 cells expressing H1.2-HaloTag (Halo). The localization of H1.2-Halo-TMR is similar to that of DAPI. **(C)** Western blotting of parental RPE-1 (left) and RPE-1 expressing H1.2-Halo (right) using an anti-H1.2 antibody. Note that the untagged H1.2 signal is reduced in RPE-1 cells expressing H1.2-Halo due to mono-allelic HaloTag insertion. The asterisk indicates a non-specific signal. **(D)** Sparse labeling of H1.2-HaloTag-TMR and oblique illumination to illuminate the thin optical layer of the nucleus for single-molecule imaging. **(E)** A representative single-molecule image of H1.2-Halo-TMR in RPE-1. Each white dot represents a single molecule of H1.2-Halo. **(F)** Single-step photobleaching of the H1.2-Halo-TMR signal. Mean intensity plots of 25 dots with standard deviation are shown after aligning the bleaching time point as 0 s. **(G)** Representative trajectories of single nucleosomes labeled with H2B-Halo-TMR (left) and H1.2-Halo-TMR (right) for 50 ms/frame. Note that H1.2 moves much more dynamically than the nucleosome. **(H)** Mean squared displacement (MSD) plots (± SD among cells) of single-H1.2 (blue) and nucleosomes (H2B-Halo; red) in living RPE-1 cells in a tracking time range from 0.05 to 1 s (n = 40 cells for each).

Oblique illumination microscopy was used to selectively illuminate a thin layer within a single nucleus (Fig. 2D) (*67, 77*). This technique, combined with sparse labeling (*67*), allowed us to visualize individual H1.2-Halo molecules as distinct dots (Fig. 2E) and track their movements, recording at 50 ms (Movies S3-S4). As a control, nucleosomes labeled with H2B-Halo in RPE-1 cells were also examined (Fig. S3A-C) (*78*). As in the case of H2B-Halo (*78*), H1.2-Halo dots exhibited single-step photobleaching (Fig. 2F), confirming that each dot represented a single H1.2 molecule labeled with Halo-TMR. The precise positions of individual dots were determined (*79, 80*) and tracked using TrackMate (*81*) to generate trajectory data. The position determination accuracy for H1.2-Halo and H2B-Halo was 9.1 nm and 8.9 nm, respectively (Fig. S2G; see Methods).

Our analysis revealed that the motion trajectories of H1.2 are highly variable, whereas nucleosome trajectories appear more constrained (Fig. 2G). We calculated the displacement (Fig. S2H) and Mean Squared Displacement (MSD) (Fig. S2I) from the trajectory data. The MSD plots showed that H1.2, on average, exhibits greater mobility than nucleosomes (Fig. 2H). This observation is consistent with previous FRAP studies (*55, 56*). We also generated a cell line with bi-allelic HaloTag insertion (Fig. S3D-G) and confirmed the H1.2-Halo in this cell behaves similarly to the one from the mono-allele (Fig. S3H-J).

### Machine learning-based H1 trajectory analysis reveals that the majority of H1 behaves like a liquid

To obtain more detailed information on H1.2 motion trajectories, we employed a machine-learning-based technique called vbSPT (*82, 83*), which can classify H1 trajectories into distinct states using a Hidden Markov Model (HMM)(Fig. 3A). Each state can be analyzed individually. Through statistical analysis based on the Richardson and Lucy (RL) algorithm (*84*), we identified three distinct states in the H1.2 trajectory data (Fig. 3B), and then categorized the data as blue, green, and red trajectories using vbSPT (Figs. 3C and S4A). The blue trajectories displayed highly constrained motion (displacement: ∼40 nm/50 ms, Fig. 3D), similar to nucleosomes (Fig. 2G). The green trajectories showed diffusive motion (∼80 nm/50 ms), while the red trajectories represented a transiently dissociated H1.2 from chromatin (∼300 nm/50 ms) (Fig. 3D).

**Fig. 3.**
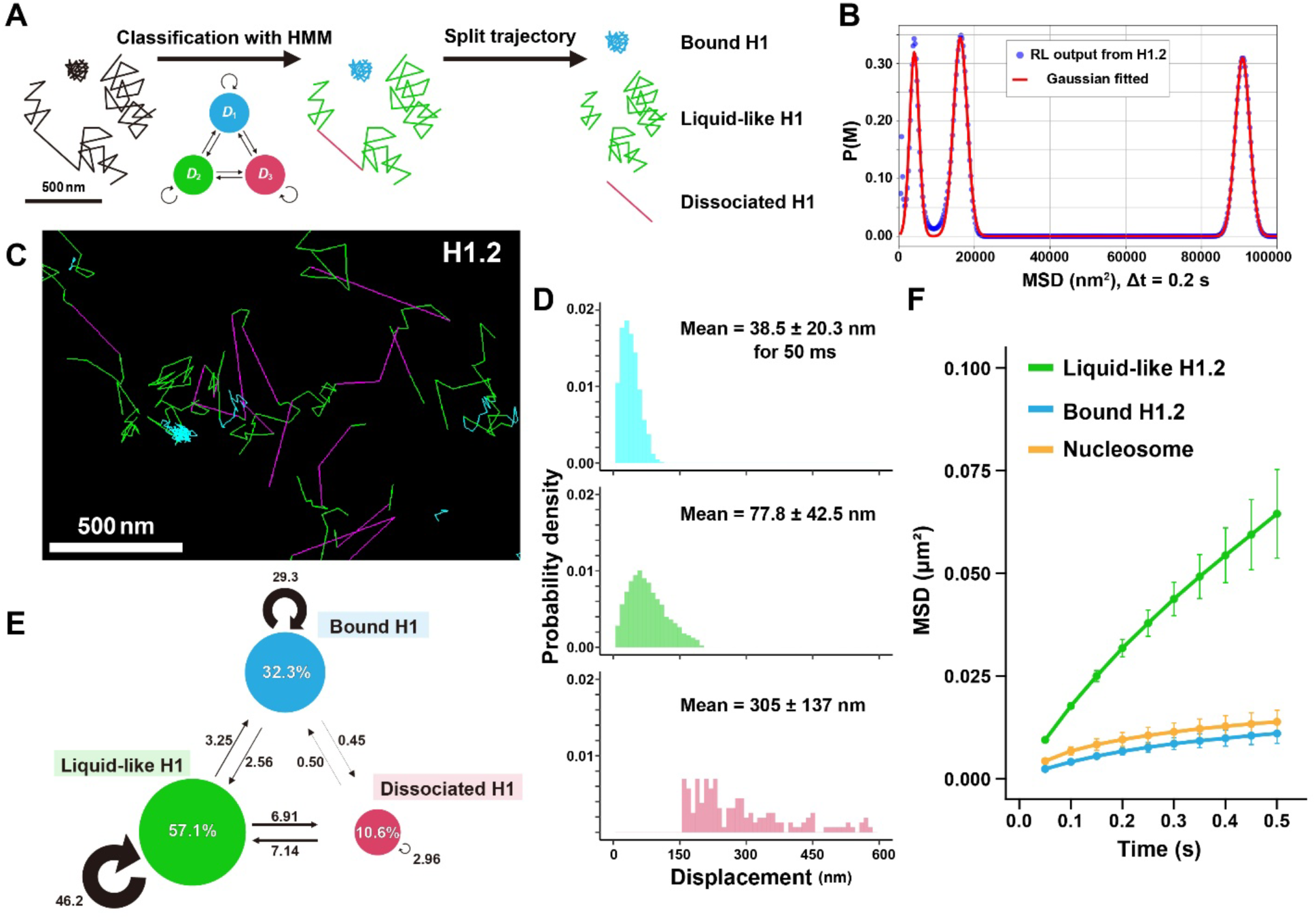
Trajectory analyses of individual H1.2-Halo molecules. **(A)** Scheme of trajectory classification by vbSPT(*82*). Based on the Hidden Markov Model (HMM), trajectories can be classified and split for further analysis. **(B)** Three prominent peaks were obtained from the MSD data of H1.2 by RL algorithm (*84*). Output from RL algorithm (blue dots) and the fitted Gaussian mixture (red line). **(C)** Classified trajectories of H1.2. “Bound H1” (cyan), constrained motion from nucleosome binding (Fig. 2G); “liquid-like H1” (green), diffusive motion; “dissociated H1” (magenta), a transient long displacement. **(D)** Displacement distribution histograms (n = 40 cells) of classified states for 50 ms. Means ± SD of displacement are at the top of each. **(E)** vbSPT results for H1.2 data (n =40 cells). *D_Bound_* : 3.54 × 10^-2^ µm^2^/s, *D_Liquid-like_* : 1.14 × 10^-1^ µm^2^/s, *D_Dissociated_* : 1.85 µm^2^/s. Arrows and numbers show the transition direction and percentage between states within 1 frame (50 ms), respectively. Note that the most frequent transition is between the liquid-like and dissociated state. **(F)** MSD plots (± SD among cells) of classified states of H1.2 and nucleosomes (H2B) in living RPE-1 cells over a tracking time range from 0.05 to 0.5 s (n = 40 cells for each).

After classification, we quantified the percentage of H1 in each state. We categorized 30% of H1, corresponding to the blue trajectories, as “bound” H1 because its motion closely resembled that of nucleosomes (Fig. 3E and 3F). Interestingly, 60% of H1, represented by the green trajectories, exhibited an almost linear increase in MSD (exponent α = 0.91) over short time scales up to ∼0.5 s (Figs. 3F and S4B), indicative of diffusive motion. This behavior led us to classify this population as “liquid-like” H1, reflecting its dynamic and fluid interaction with chromatin. The remaining 10% was classified as “dissociated” H1. These findings suggest that the majority of H1 behaves in a liquid-like manner. We therefore focused our subsequent analysis on this liquid-like H1 population.

To exclude the possibility that the three states of H1 motion (bound, liquid-like, and dissociated) are specific to H1.2-Halo, we created RPE-1 cells stably expressing H1.0-Halo using a similar strategy as for H1.2-Halo (Fig. S4C-F), and then investigated the behavior of the H1.0-Halo molecules in live RPE-1 cells (Fig. S4G). Consistent with the results for H1.2-Halo, we identified three distinct states in the H1.0 trajectory data (Fig. S4H-I): “bound” (27.7%), “liquid-like” (58.0%), and “dissociated” (14.3%). We concluded that the three states of H1 motion are not variant-specific and that most H1 interacting with chromatin behaves like a liquid.

### H1 diffuses within chromatin domains in a liquid-like manner

We sought to determine whether H1 diffuses like a liquid inside or outside chromatin domains (Fig. 4A). To distinguish between these locations, we visualized chromatin domains in living cells using PALM (photoactivated localization microscopy) super-resolution imaging (*8, 85*) of nucleosomes and combined this with single H1 imaging. We labeled nucleosomes and H1.2 in different colors: H2B-PAmCherry (*8*) for nucleosomes and H1.2-Halo-PA-JF646 (*86*) for H1.2 (Figs. 4B and S5A). Simultaneous observations of nucleosomes and H1 dots were achieved using a beam splitter system (Figs. 4B-C and S5B) (*12, 87*). Chromatin domains were reconstructed from the PALM data, and we collected over 100,000 H1.2 trajectories from a single cell, which were classified by vbSPT into bound, liquid-like, and dissociated states (Movies S5-S6).

**Fig. 4.**
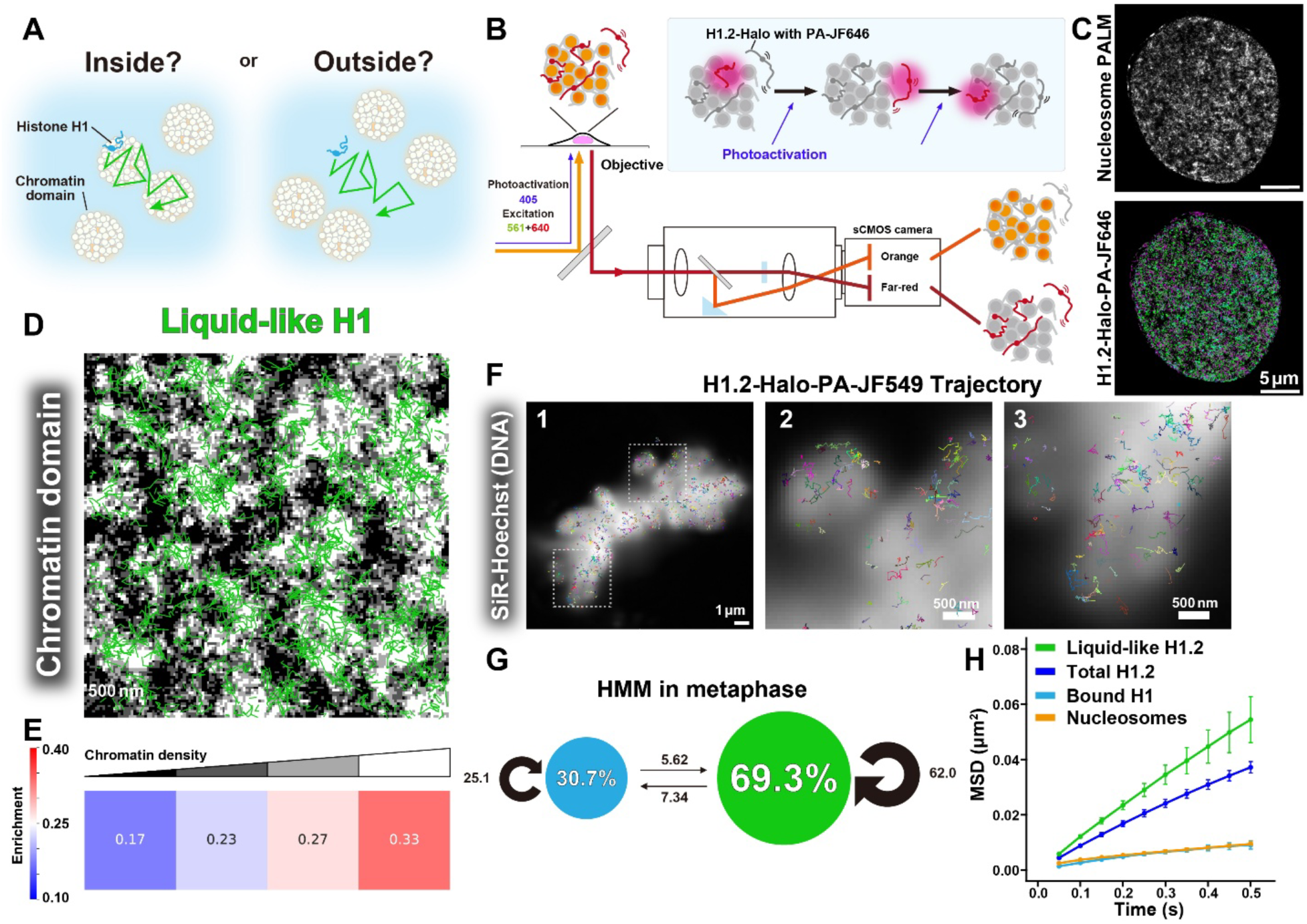
Liquid-like behavior of H1.2 within the chromatin domain and mitotic chromosome. **(A)** Question scheme: Does H1 diffuse like a liquid inside or outside chromatin domains? **(B)** Schematic for dual-color imaging of RPE-1 cells expressing H1.2-Halo and H2B-PAmCherry with a beam splitter system. Box, photoactivation scheme of H1.2-Halo-PA-JF646. The images of H1.2-Halo-PA-JF646 and H2B-PAmCherry were simultaneously acquired with a single sCMOS camera (left half, orange color; right half, far-red color). PA-JF646 and PAmCherry were continuously activated using a very weak 405 nm laser. After PALM reconstruction of H2B-PAmCherry signals, chromatin domains and motion of H1 were visualized at the same time with nm-scale resolution. **(C)** Nucleosome PALM image constructed from H2B-PAmCherry sequential images (50 ms/frame, obtained over 50 s) in an RPE-1 cell (upper), and classified trajectories of H1.2 (bottom) in the same cell. **(D)** Overlap of the PALM image of nucleosomes and trajectories of liquid-like H1. Note that liquid-like H1 shows clear overlap with chromatin domain structures. **(E)** PALM images were visualized with four classes of nucleosome density. Enrichment of liquid-like H1 trajectories in dense chromatin regions. A value of 0.25 represents a random distribution of trajectories. **(F)** Overlap of SiR-Hoechst (DNA) signal and trajectories of H1 in metaphase chromosomes (panel 1). Individual trajectories were randomly colored. Panels 2-3 are enlarged images from the squared regions in the panel 1. H1 shows diffusive movement within mitotic chromosomes. **(G)** vbSPT results for H1.2 data in metaphase (n = 10 cells). *D_Bound_* : 4.63 × 10^-2^ µm^2^/s, *D_Liquid-like_* : 1.25 × 10^-1^ µm^2^/s. The trajectories consist only of “bound H1” and “liquid-like H1”. **(H)** MSD plots (± SD among cells) of single H1.2 and nucleosomes based on the classified states of H1.2 in living RPE-1 cells in metaphase over a tracking time range from 0.05 to 0.5 s (n = 10 cells for each).

We overlaid the liquid-like H1.2 trajectories onto the chromatin domain image (Figs. 4D and S5C), revealing a clear overlap between the domains and H1 trajectories. Quantitative analysis showed that areas with higher chromatin density (appearing whiter) were more enriched with liquid-like H1.2 (Figs. 4E and S5C). These findings suggest that H1 moves within the chromatin domain in a liquid-like manner, supporting our model that H1 acts as a liquid-like “glue” for chromatin (Fig. 1F).

Our finding that chromatin domains were more enriched with liquid-like H1.2 was further corroborated by observations of H1.2 in mitotic chromosomes (Fig. 4F). These large, condensed chromatin structures with diameters of approximately 500-700 nm (*43*) provided a clearer view of H1 behavior. Dual color imaging of single H1.2 molecule and mitotic chromosomes stained by SiR-Hoechst (DNA), clearly visualized that H1 moves within the mitotic chromosome (Figs. 4F and S5D). Using the RL algorithm (*84*), we identified two distinct states in the H1.2 trajectory data from mitotic chromosomes (Fig. S5E) and classified them with vbSPT (Fig. 4G). This analysis further affirmed our earlier results by showing that 30% of H1 was in the bound state, 70% exhibited liquid-like behavior (Fig. 4G-H), and no dissociated H1 was detected.

### Multiscale chromatin modeling supports a liquid-like glue property of H1

To test our model that H1 acts as a liquid-like glue, we examined the behavior of H1.2 within a 108-nucleosome domain (one H1.2 molecule per nucleosome) via unbiased MD simulations using our multiscale model (Fig. 1A, right; Figs. 5A and S6A; Movie S7)(*72*). The results demonstrated that H1.2 diffuses throughout the domain, forming transient multivalent interactions by simultaneously engaging multiple nucleosomes (Fig. 5B-E; Fig. S6B). Interestingly, the region-specific contact map revealed that the nucleosome dyad was less contacted region for H1.2 (Fig. 5C), consistent with its liquid-like behavior (Fig. 4D). Notably, reducing the number of H1.2 molecules by 75% within the domain (Fig. 5F, left) led to a partial decompaction of the chromatin structure (Fig. 5F, right; Fig. S6C). These findings further support our model that H1 acts as a liquid-like glue for chromatin.

**Fig. 5.**
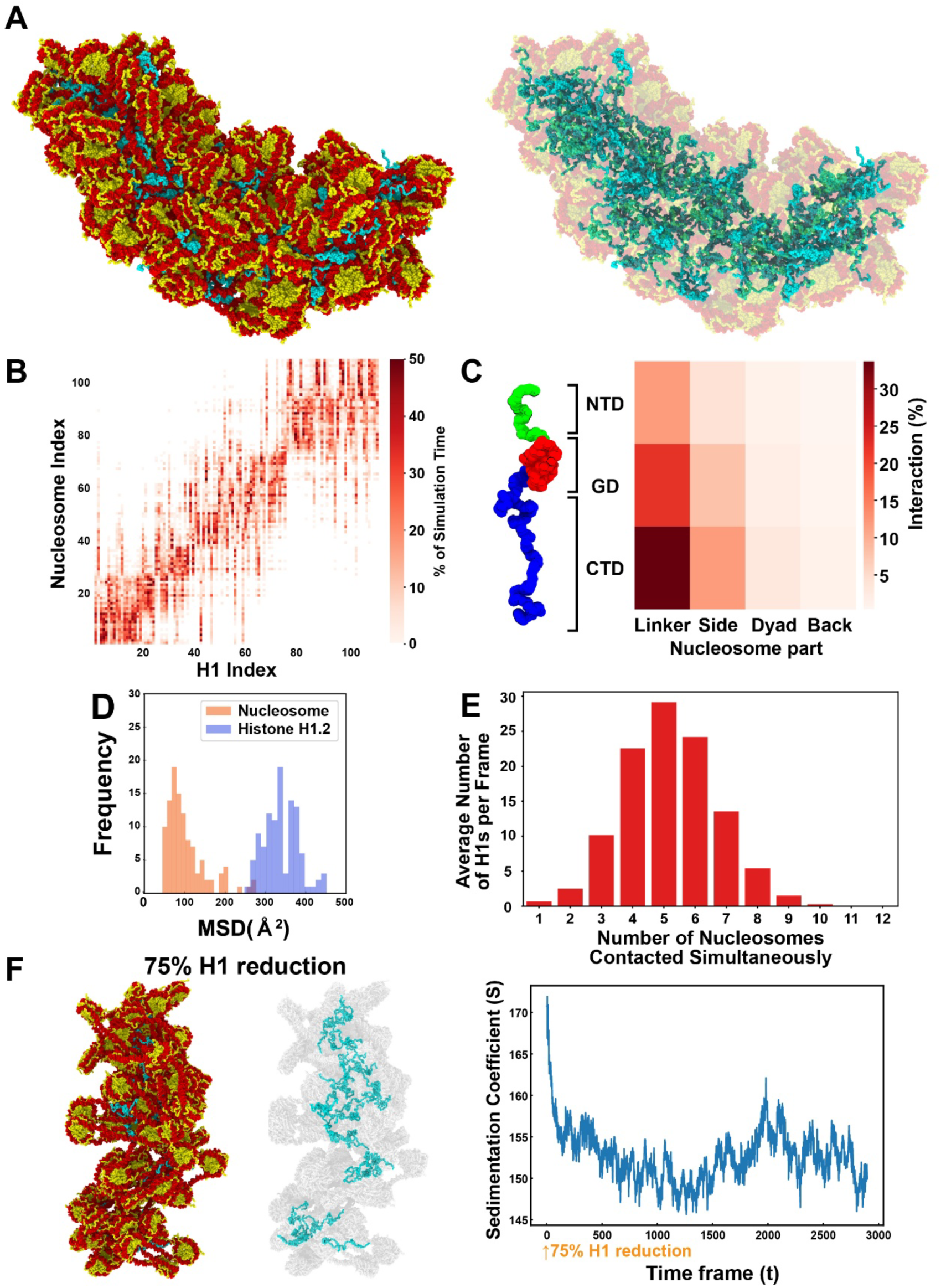
Computational simulation of 108 nucleosomes with H1.2 and a glue activity of H1.2. **(A)** The final configuration of an irregular cluster of 108 nucleosomes (195 bp NRL) with 108 H1.2 molecules. (also see Movie S7). **(B)** Contact map of 108 nucleosomes and 108 H1.2 molecules. Diagonal indicate where H1.2 is located at the beginning of the simulation. Note that H1.2 diffuses among several nucleosomes. **(C)** Region-specific contact map of H1.2 and nucleosome within a 108-nucleosome cluster. **(D)** Histograms for the MSD ( Δ t = 1 ns) for nucleosomes and H1.2 within a 108-nucleosome cluster. H1.2 has greater mobility than nucleosomes. **(E)** Histogram of the number of nucleosomes contacted simultaneously by an H1.2 during the 108-nucleosome cluster simulation in Fig. 5A-C. **(F)** When 75% of H1.2 was removed from the 108-nucleosome cluster, the cluster became decondensed (left, full configuration after reduction; right, only H1.2 are shown). (Right) The sedimentation coefficient (S) plot over time is also shown. The sedimentation coefficient drastically decreased after H1 reduction.

### H1.2 is actively involved in chromatin compaction in human cells

Finally, we investigated whether H1.2, among the H1 variants, plays a direct role in chromatin compaction in human cells. To this end, we used the auxin-inducible degron (AID) system 2 (*71*) for rapid H1.2 depletion, combined with PALM (photoactivated localization microscopy) super-resolution imaging of nucleosomes labeled with H2B-Halo. Using CRISPR-Cas9 genome editing, we inserted a cassette encoding mini-AID (mAID) and the fluorescent protein mClover at the C-terminal site of the endogenous H1.2 gene locus (Fig. 6A) in human colon adenocarcinoma HCT116 cells expressing OsTIR1(F74G) (for details, see Methods)(*71*) and H2B-Halo (Fig. S7A)(*78*). Proper insertion of the tag sequence and expression of mAID-mClover were confirmed by PCR (Fig. S7B-C) and western blotting (Fig. 6B). The distribution of H1.2-mAID-mClover (mAC) in HCT116 cells resembled the pattern observed with DAPI staining, including DAPI-dense heterochromatin (Pearson’s correlation = 0.86, Figs. 6C and S7D), indicating a proper genome-wide localization. Stepwise-salt washing (Fig. S7E) also confirmed appropriate chromatin association of H1.2-mAC. In these cells, H1.2 was effectively depleted within 3 hours after adding the auxin analog 5Ph-IAA (Figs. 6B, 6D-E, and S7F).

**Fig. 6.**
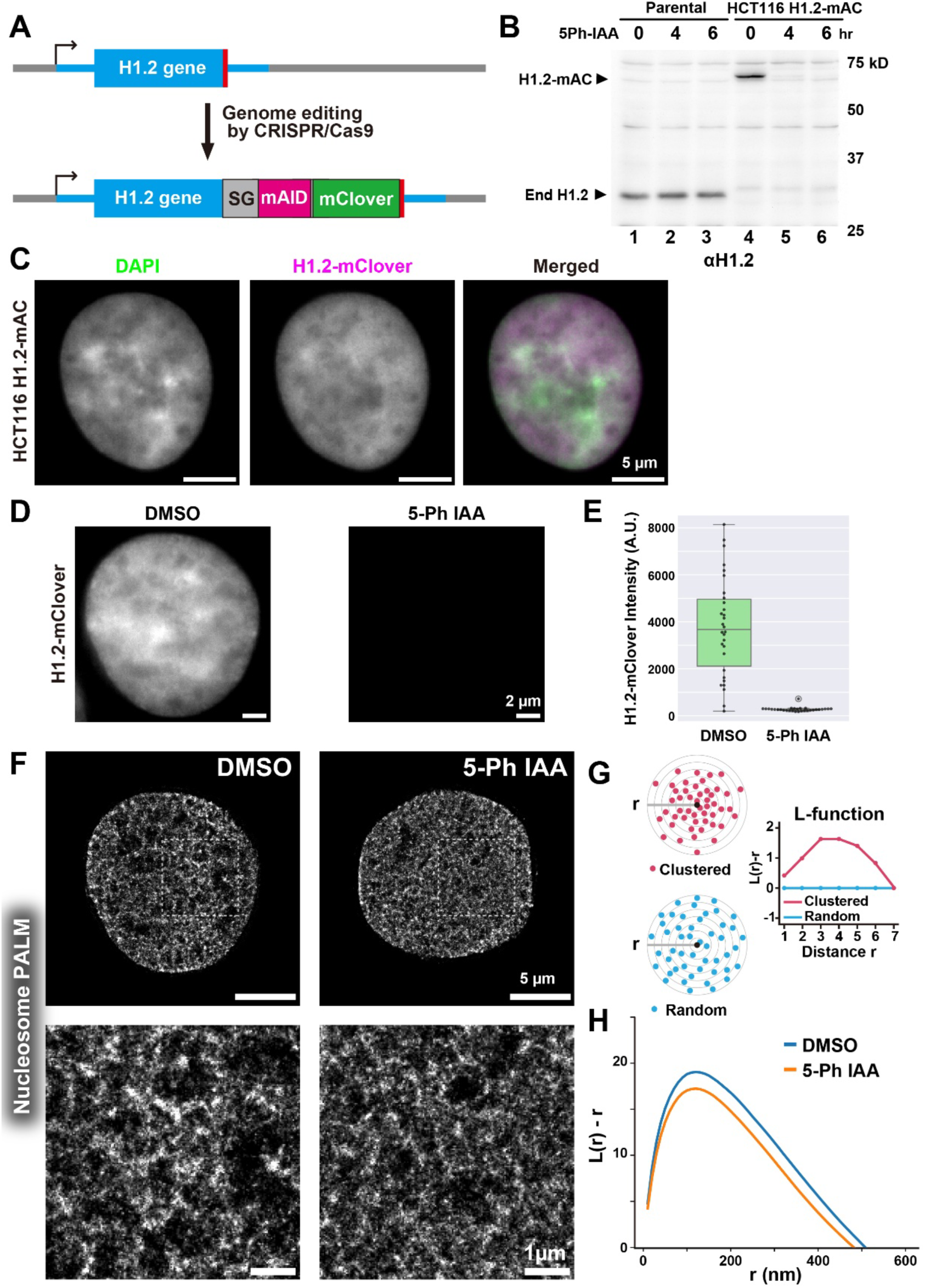
Rapid-depletion of H1.2 protein in HCT116 decondensed chromatin domains. **(A)** Schematic of CRISPR/Cas9-mediated genome editing for inserting mAID and mClover (mAC) at the C-terminus of the H1.2 gene locus in HCT116 cells expressing OsTIR1(F74G). **(B)** Rapid-depletion of H1.2 upon 5Ph-IAA addition. Western blotting of H1.2 for parental HCT116 and HCT116 cells expressing H1.2-mAC using an anti-H1.2 antibody. Cells were treated with 1 µM 5Ph-IAA for the indicated time. **(C)** Representative images of FA-fixed HCT116 cells expressing H1.2-mAC. The localization of H1.2-mAC is similar to that of DAPI. **(D)** Representative images of H1.2-mClover signal in FA-fixed HCT116 cells expressing H1.2-mAC with 0.01% DMSO (left) or 1 µM 5-Ph IAA for 3 hours (right). **(E)** Quantification of depletion efficiency based on H1.2-mClover intensity in FA-fixed HCT116 cells expressing H1.2-mAC, treated with either DMSO (left) or 5-Ph IAA (right) (n = 30 cells per condition). **(F)** PALM images of nucleosomes labeled with H2B-Halo-PA-JF646 in FA-fixed HCT116 cells expressing H1.2-mAC, treated with either DMSO (left) or 5-Ph IAA (right). Enlarged views of the squared regions in the upper panels are shown at the bottom. Notably, the H1.2-depleted cell exhibits a more dispersed chromatin domain structure. **(G)** A simplified schematic of L-function analysis. Shown are clustered (red spheres, top left) and random (blue spheres, bottom left) particles surrounding the origin point (black sphere). L-function plots for a random pattern (blue) are approximately 0. **(H)** L-function plot of chromatin in DMSO-treated (control, blue) and 5-Ph IAA-treated (H1-depleted, orange) cells (n = 30 cells per condition). L-function values decreased with H1 depletion, indicating chromatin domain decondensation.

To perform PALM imaging of nucleosomes labeled with H2B-Halo-PA-JF646 (*86*), we fixed cells treated with or without 5Ph-IAA using formaldehyde, then reconstructed PALM images of the nucleosomes. The reconstructed images showed chromatin decondensation following rapid H1.2 depletion (Fig. 6F). This finding was further supported by a decrease in the L-function plot (L(r)-r versus r), which estimates the size and compaction state of nucleosome clusters (or chromatin domains) (Figs. 6G-H, and S7G) (*8, 88*). These results indicate that H1.2 is involved in chromatin domain condensation, consistent with our modeling results (Figs. 5F and S6C). Furthermore, they also suggest that tagged H1.2 (with mAID and mClover) retains its functionality.

## Discussion

Using single-molecule imaging, we quantitatively tracked the motions of linker histone H1 and nucleosomes in interphase chromatin and mitotic chromosomes with unprecedented spatiotemporal resolution (10 nm/millisecond scale). By leveraging machine-learning-based motion trajectory analysis (*82, 83*), we uncovered three distinct states of H1 behavior in chromatin. This analysis provides a mechanistic understanding of H1 binding modes in live cells.

Surprisingly, 60-70% of H1 binds to nucleosomes dynamically and exhibits liquid-like behavior within the chromatin domain and mitotic chromosomes. Since X-ray crystallography and cryo-EM studies typically require static and regular structures, this flexible and dynamic state of H1 might not have been well captured. Our findings indicate that the liquid-like state is the primary binding mode of H1 in cellular chromatin (Fig. 7), also consistent with recent evidence that the 30-nm fiber is not the fundamental chromatin structure in living cells(*25, 51–53*). According to the modeling results (Fig. 5E), H1 can interact transiently with several nucleosomes simultaneously, enabling the formation of a liquid network of weak multivalent interactions between H1 and nucleosomes. These findings suggest that H1 functions as a liquid-like “glue” for the chromatin domain, with its positively charged long tails weakly adhering to DNA and acting as a weak glue (Fig. 7).

**Fig. 7.**
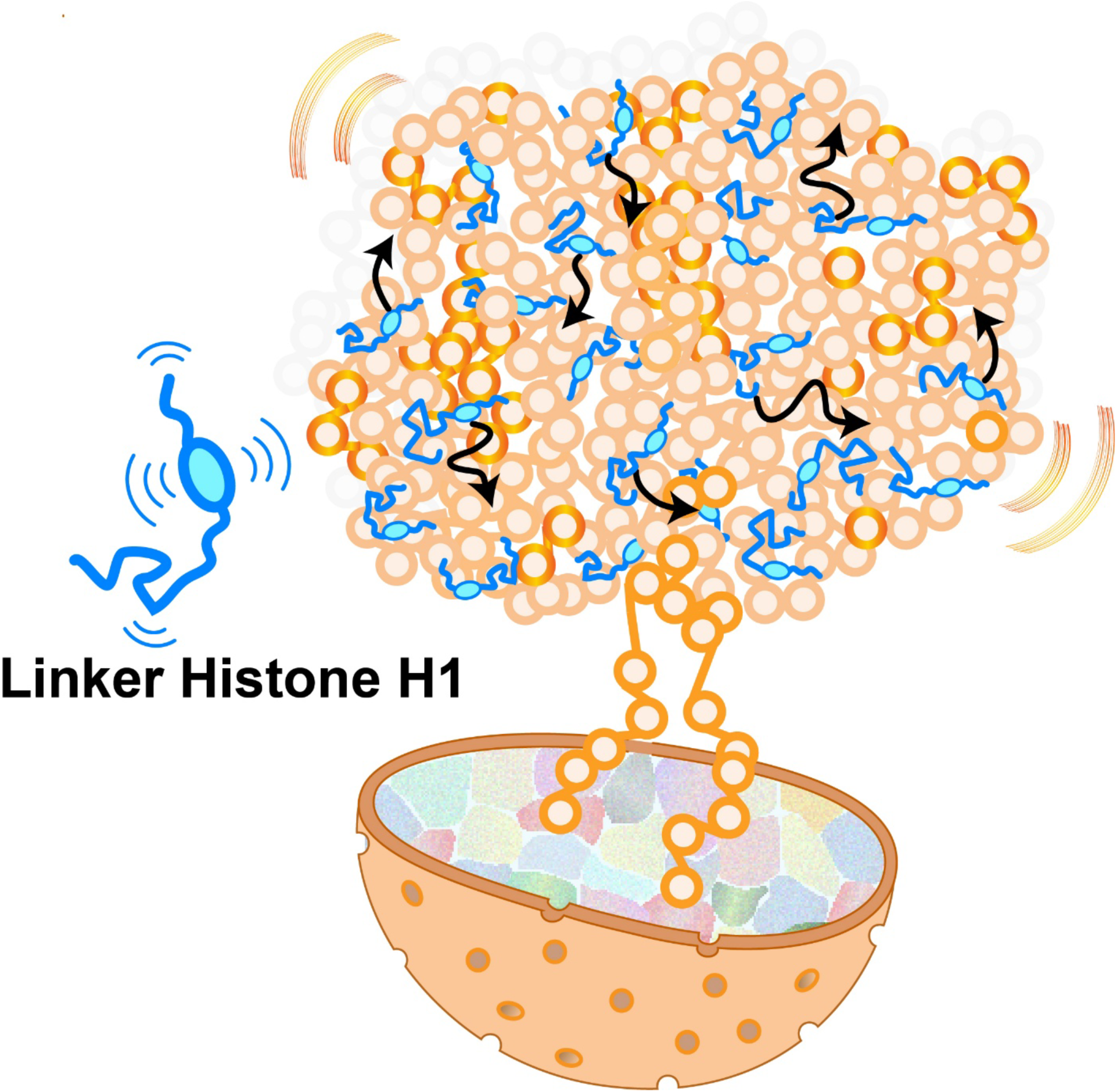
Model figure. Linker histone H1 functions as liquid-like “glue” for chromatin condensation. H1 dynamically interacts with nucleosomes via a positively charged CTD with multivalent interactions. Such liquid-like “glue” activity can condense chromatin while keeping chromatin fluid and accessible. Note: This model is highly simplified—many more H1 molecules exist and move dynamically within chromatin in living cells.

Our liquid-like H1 model is compatible with the findings of Turner et al. (*89*). Turner et al. reported that the CTDs of chicken histone H1 remain disordered and dynamic when bound to short double-stranded DNA (dsDNA), forming a liquid-like condensate (H1 condensate was also reported in (*90*)). This report suggests that the CTDs of linker histones can function as a liquid-like “glue” through dynamic linker histone–nucleosome interactions, which facilitate crosslinking of linker DNA segments (*89, 91*). Indeed, when a mutant H1, which has a CTD lacking half of its positive charge, was ectopically expressed, the fraction of “liquid-like” H1 drastically decreased (61.5% > 24.8% in Fig. S8), suggesting that the CTD’s disorder and high positive charge is crucial for the liquid-like “glue” property of H1.

Our observation also aligns with previous reports using FRAP experiments (*55–58, 92*). Notably, our direct H1 measurements (Fig. 3E) are in good agreement with the predicted fractions of H1 binding modes (slow, 28 ± 6%; fast, 71 ± 5%; free, 0.4 ± 0.1%), which were calculated from FRAP results using a reaction-diffusion model (*57*).

It is also intriguing to discuss the theoretical aspects of the liquid-like glue property of H1. We calculated the stable state of H1 using a mean-field model, similar to the polymer model of Flory and Huggins (*93*). This model accounts for the multivalent interactions between H1 and nucleosomes, and the entropy gain due to the dynamic binding of mobile H1 molecules. The results predicted that the majority of H1 would adopt a mobile state in nucleosome-dense conditions (Fig. S9), indicating condensed chromatin is more stable with mobile H1 due to its ability to form inter-nucleosomal multivalent interactions. Notably, this theoretical prediction is consistent with our experimental observations and model, which show that ∼60-70% of H1 behaves like a liquid within chromatin domains and mitotic chromosomes, functioning as a dynamic “glue” (Figs. 3E and 4G).

In contrast, we found that approximately 30% of H1 stably binds to nucleosomes, which may correspond to “dyad” binding in the classical model. This proportion remained consistent in both interphase chromatin and mitotic chromosomes. Therefore, our study suggests that stable “dyad” binding is not the primary mode of H1 interactions in live cells and occurs mainly as a transient event. “Dyad” binding can condense chromatin by restricting the orientation and flexibility of linker DNA. This raises intriguing questions: Are there specific functions associated with this stable binding mode in cellular chromatin? Could it be genome sequence-specific? Is there a relationship with particular histone variants or modifications? Exploring these possibilities will provide deeper insight into the functional roles of H1 binding in chromatin organization.

Our findings suggest that H1 acts as a liquid glue, compacting chromatin while maintaining DNA accessibility to proteins, as the nucleosomes within chromatin domains and mitotic chromosomes “glued” together by H1 remain highly mobile. This dynamical mode of compaction may offer several advantages for template-directed biological processes during interphase, such as RNA transcription and DNA replication/repair/recombination. For example, in transcriptional regulation, the liquid-like movement of H1 may facilitate the diffusion of transcription factors and complexes into targets within condensed chromatin domains. Even in mitosis, cells must maintain proper accessibility for large factors such as condensins and topoisomerase IIα, which are involved in chromosome assembly (*94, 95*). The dynamic, liquid-like behavior of H1 likely enables these proteins to efficiently penetrate into condensed chromatin (*64, 87, 96*). The liquid-like H1 binding mode may thus contribute to the assembly of mitotic chromosomes.

## Methods

### Multiscale MD simulations of chromatin arrays

The model explicitly represents each amino acid in the histone proteins with one bead centered on its *C*_α_. Beads corresponding to lysine, arginine, aspartic acid, glutamic acid, and histidine carry the total charge of their atomistic counterparts at pH = 7. The relative hydrophobicities and diameters of all amino acid beads were derived from atomistic simulations and experimental data as described in (*72*).

To preserve the secondary structure of the histones within the histone core we used an elastic network model defined by our atomistic MD simulations of mono-nucleosomes (*70*), which were based on the 1KX5 crystal structure (*97*). In contrast, the histone tails were modelled as fully flexible polymers, with bonds between consecutive residues maintained via a stiff harmonic potential, and no energetic penalty considered for bending or torsion.

DNA was modelled at a resolution of one ellipsoid per base pair and one point charge per phosphate group (*72*). The sequence-dependent mechanical properties of DNA were implemented using a modified version of the Rigid Base Pair model (*72, 98*), which approximates inter-base pair step deformations with a six-dimensional harmonic potential (shift, slide, rise, roll, tilt, and twist) using parameters derived from large-scale atomistic simulations of free DNA strands (*99*).

Electrostatic interactions between all non-bonded beads were approximated using the Debye-Hückel model (*100*), while non-ionic associations were modeled using a Lennard-Jones potential (*101*). Parameters for amino acid pair interactions were taken from the Kim-Hummer model (*102*), and amino acid–DNA parameters were derived from atomistic simulations of nucleosomes (*70*). Further details of all model parameters and energy functions are available in (*72*).

For the 12-nucleosome array simulations, we tested two regular nucleosome repeat lengths: 165 bp and 195 bp. The initial configurations included one H1.2 molecule bound to each nucleosome in its dyad position. All the simulations were performed for the human H1.2 protein (Uniprot Consortium ID: P16403). We defined the N-terminal domain as residues 1-38, and the C-terminal domain as residues 109-213. We used Modeller to create a homology model of the globular domain (GD) of H1.2 with H5 (PDB 4QLC) as the template. In all the simulations, H1 proteins can bind/unbind dynamically from the dyad position and diffuse around chromatin freely.

For efficient sampling of chromatin configurations, we performed temperature-replica exchange molecular dynamics simulations (T-REMD) at 0.15 M NaCl (Debye length of 8 Å). The number of replicas and spacing were optimized to ensure acceptance probabilities exceeding 30% between neighboring replicas. We required 64 replicas to span temperatures between 300K and 600K. These simulations were performed in the canonical ensemble (NVT) using the Langevin thermostat (damping time of 10,000 fs) and the velocity Verlet integrator implemented in LAMMPS (*103*). The simulations were run for approximately 100 million steps, using a timestep of 40 fs. T-REMD exchanges were attempted every 1,000 steps and simulation snapshots were recorded every 100,000 steps.

For the 108-nucleosome array simulations, we built the system using 9 pre-equilibrated configurations of 12-nucleosome arrays. This is an unbiased simulation run at 0.15 M NaCl (Debye length of 8 Å) and 300K for illustration purposes.

For analysis, we computed sedimentation coefficients using the HullRad method (*104*), which uses a convex hull model to estimate the hydrodynamic volume of chromatin. Specifically, the hydrodynamic volume is approximated by constructing the smallest convex envelope around chromatin while accounting approximately for hydration effects and solvent interactions. The HullRad method is particularly suited for flexible and irregular structures like chromatin.

### Plasmid construction

Construction of the pX330 CRISPR-Cas9 plasmid (#42230, Addgene) expressing guide RNA for H1.2 or H1.0 target sites and the donor plasmids were performed as follows. Gene-specific guide RNA sequences for H1.2 or H1.0 were designed using the CRISPR design website and inserted into the pX330 BbsI cloning site (*73*). The guide RNA sequences were as follows: For H1.2, 5’-CACCGGGTTGTCAAGCCTAAGAAGG–3’ and 5’-AAACCCTTCTTAGGCTTGACAACCC-3’.

For H1.0, 5’-CACCGCTTGCCGGCCCTCTTGGCAC–3’ and 5’-AAACGTGCCAAGAGGGCCGGCAAGC-3’.

For the donor plasmids of H1.2-HaloTag and H1.0-HaloTag, the left and right homologous arms were PCR amplified using KOD FX (KFX-101, Toyobo) from RPE-1 genomic DNA, which was extracted using a Wizard Genomic DNA Purification kit (A1120, Promega). The primer sequences were as follows: For H1.2 of left arm, 5’-AGATCCACCTCCACCTTTCTTCTTGGGCGCTGCCTTCTTAGGCTT-3’ and 5’-GCGGCCGCGGGAATTCCTCCTGCCGCTCCCGCTGC-3’; H1.2 of right arm, 5’-GAGATTTCCGGTTAATAGGCGAACGCCTACTTCTA-3’ and 5’ CTCCCATATGGTCGACGGACAAAATGGCTGGCTAAAAGT–3’. For H1.0 of left arm, 5’-GCGGCCGCGGGAATTCTCCACAGACCACCCCAAGTA-3’ and 5’-AGATCCACCTCCACCCTTCTTCTTGCCGGCTCTCTTGGCAC-3’; H1.0 of right arm, 5’-GAGATTTCCGGTTAATGACAATGAAGTCTTTTCTT-3’ and 5’-CTCCCATATGGTCGACTCGGGAGGTTTTAAGTGGCC–3’.

The HaloTag sequence was PCR amplified from the pFC14A HaloTag CMV Flexi Vector (G965A, Promega) using the following primers: 5’– GGTGGAGGTGGATCTGGTGGAGGTGGATCTGGTGGCGGCGGTTCAGGATCCGAAATCGGTACTGGCTTTC–3’ and 5’–TTAA CCGGAAATCTCCAGAG–3’. The homologous arms and HaloTag fragment were joined using standard overlapping PCR and inserted between the EcoRI and SalI sites of the pGEM-T (Easy) vector (A1360, Promega) using In-Fusion (639648, Clontech).

To construct the donor plasmid for H1.2-mAID-mClover (mAC), sequences except HaloTag were amplified from the H1.2-HaloTag donor plasmid using the following primers: 5’-TGAACCGCCGCCACCAGATC-3’ and 5’-GCGAACGCCTACTTCTAAAACC-3’. mAID-mClover was amplified from mAC-POLR2A donor (Hygro)(#124496, Addgene) with 5’–GGTGGCGGCGGTTCAAAGGAGAAGAGTGCTTGTCC –3’ and 5’–GAAGTAGGCGTTCGCTTACTTGTACCAAGGCCTTCCGTCCATGC–3’. Both fragments were joined using In-Fusion.

For vector construction to transiently express wildtype H1.2-HaloTag (H1.2-WT), fragments were amplified from a donor plasmid of H1.2-Halo using the following primers: 5′-CAGCGGGAGCGGCAGGAGCAGTCTCGGACATGGTGGCGTTAATTAACCTTAAGTTTACGAGGG-3′ and 5′-CTCTGGAGATTTCCGGTTAATAGACCGGTGCGGCCGCAATCGATCGCC-3′. The amplified H1.2-Halo fragments were joined together using standard overlapping PCR and inserted into the PacI and AgeI sites of pAAVS1-NDi-CRISPRi (#73498, Addgene) using the XE cocktail (*105*). The backbone part of the H1.2-WT vector was amplified using 5’‒GGATCCGAAATCGGTACTGGCTTTC‒3’ and 5’‒CTCATCACCAAGGCTGTGGCCG‒3’.

Then, the synthesized H1.2-Khalf CDS (gBlocks, IDT) and the backbone were ligated using the XE cocktail.

### Establishment of stable cell lines and cell culture

To establish RPE-1 cells that stably express H1.2-HaloTag from the endogenous locus, RPE-1 cells at 80% confluence in a 6-well dish were cotransfected with 750 ng each of pX330 (with a target gene-specific guide sequence inserted) and the H1.2-Halo donor plasmid using the Neon electroporation transfection system (ThermoFisher, MPK1025). Twenty-four hours after transfection, transformants in the 6-well dish were transferred to a 10-cm dish and expanded until they reached 5 x 10^6^ cells/mL. To select transfected cells from untransfected cells, a cell sorter (SONY, SH800S) was used. Before sorting, cells were incubated in a medium containing 50 nM TMR-HaloTag ligand (8251, Promega) for 16 hours. FACS profiles and Gates used to collect H1.2-HaloTag-TMR positives are also indicated in Fig. S2A. The number of cells collected per total number of cells analyzed is also indicated for each run. Cells were collected into 2 mL of ice-cooled collection medium (DMEM consisting of 50% fresh and 50% conditioned medium (harvested from log phase growth cells and 0.2 µm filtered)). FBS and penicillin streptomycin were added to final concentrations of 20% and 2.5 µg/mL, respectively. Cells were expanded and colonies were isolated. Proper insertions were confirmed by PCR with the following primers: 5’‒GGCCCCTGTAAAGAAGAAGG‒3’ and 5’‒GGACAAAATGGCTGGCTAAA‒3’.

For RPE-1 cells that stably express H1.0-HaloTag from the endogenous locus, we used a similar strategy for H1.2-HaloTag HCT116 cells. Proper insertions were confirmed by PCR with the following primers: 5’‒AGGAAAAGCAGCGACTCCTC‒3’ and 5’‒CTGCTCCCTACCACCTCTCT‒3’.

To further express H2B-PAmCherry in the obtained RPE-1 cells, pPB-PGKneo-EF1α-H2B-PAmCherry-BGH-polyA (*8*) and pCMV-hyPBase (provided by the Sanger Institute with a materials transfer agreement) were co-transfected to the RPE-1 expressing H1.2-Halo with the Effectene Transfection Reagent kit (301425; QIAGEN). The transformants were then selected using 600 μg/mL G418.

To create HCT116 H1.2-mAC cells, HCT116 with CMV-OsTIR1-F74G (*71*) were cotransfected with 750 ng each of pX330 (with a target gene-specific guide sequence inserted) and the H1.2-mAC donor plasmid using the Neon electroporation transfection system (ThermoFisher, MPK1025). The transformants were enriched using a cell sorter and colonies were isolated. Proper insertions were confirmed by PCR with the following primers: 5’‒GGCCCCTGTAAAGAAGAAGG‒3’ and 5’‒GGACAAAATGGCTGGCTAAA‒3’. To further express H2B-Halo in the obtained HCT116 H1.2-mAC cells, pPB-CAG-IB-H2B-HaloTag (*78*) and pCMV-hyPBase were co-transfected. The transformants were then selected with 10 μg/mL blasticidin S (029–18701; Wako; pPB-CAG-IB-H2B-HaloTag).

RPE-1 cells were cultured in Dulbecco’s Modified Eagle’s Medium (DMEM; D5796; Sigma-Aldrich) supplemented with 10% fetal bovine serum (FBS; F7524, Sigma) at 37℃ under 5% CO_2_. HCT116 cells were cultured in McCoy’s 5A medium (16600-082; Gibco) supplemented with 10% fetal bovine serum (FBS; F7524, Sigma) at 37℃ under 5% CO_2_.

### Biochemical fractionation of nuclei from cells expressing H1.2-HaloTag

Nuclei were isolated from RPE-1 cells expressing endogenous H1.2-HaloTag as described previously (*106*). Briefly, collected cells were suspended in nuclei isolation buffer (3.75 mM Tris-HCl (pH 7.5), 20 mM KCl, 0.5 mM EDTA, 0.05 mM spermine, 0.125 mM spermidine, aprotinin (1 µg/mL) (T010A; TaKaRa), and 0.1 mM phenylmethylsulfonyl fluoride (PMSF) (P7626-1G; Sigma-Aldrich)) and centrifuged at 1936 × g for 7 min at room temperature. The cell pellets were resuspended in nuclei isolation buffer and again centrifuged at 1936 × g for 7 min at room temperature. Subsequent steps were performed at 4°C unless otherwise noted. Cell pellets were resuspended in nuclei isolation buffer containing 0.025% Empigen (45165-50ML, Sigma-Aldrich) (nuclei isolation buffer+) and homogenized immediately with 10 downward strokes of a tight Dounce pestle (357546; Wheaton). The cell lysates were centrifuged at 4336 × g for 5 min. The nuclear pellets were washed in nuclei isolation buffer+. The nuclei were incubated on ice for 15 min in the following buffers containing various concentrations of salt: HE (10 mM HEPES-NaOH (pH 7.5), 1 mM EDTA, and 0.1 mM PMSF), HE + 100 mM NaCl, HE + 200 mM NaCl, HE + 300 mM NaCl, and HE + 500 mM. After each buffer incubation with increasing concentrations of salt, centrifugation was performed to separate the nuclear suspensions into supernatant and pellet fractions. The proteins in the supernatant fractions were precipitated by using 17% trichloroacetic acid (208-08081; Wako) and cold acetone. Both pellets were suspended in a Laemmli sample buffer and subjected to 12.5% SDS-PAGE, followed by Coomassie brilliant blue (031-17922; Wako) staining and western blotting using rabbit anti-H1.2 (19649-1-AP; Proteintech) and rabbit anti-HaloTag (G9281; Promega) antibodies to confirm H1.2-HaloTag expression.

### Bulk Halo-tag labeling and fixed cell imaging

H2B-HaloTag or H1.2/H1.0-HaloTag expressing cells were grown on poly-L-lysine coated (P1524-500MG, Sigma-Aldrich) coverslips (C018001, Matsunami) and labeled with 40 nM HaloTag TMR ligand (G8251, Promega) overnight. The cells were fixed with 1.85% formaldehyde (FA) (064-00406, Wako) at room temperature for 15 min, permeabilized with 0.5% Triton X-100 (T-9284, Sigma-Aldrich) for 5 min, and stained with 4’,6-diamidino-2-phenylindole (DAPI; 0.5 µg/mL) (10236276001, Roche) for 5 min, followed by PPDI (20 mM Hepes (pH 7.4), 1 mM MgCl_2_, 100 mM KCl, 78% glycerol, and paraphenylene diamine (1 mg/mL) (695106-1G, Sigma-Aldrich)) mounting. Z-stack images with 0.2 µm thickness were acquired with a DeltaVision system (Applied Precision) with a UPlanApo 60× 1.40 NA objective lens (Olympus). The z-stack images were deconvolved with softWoRx (Applied Precision) and presented as maximum intensity projections of 5 sections.

### HaloTag labeling for single molecule imaging

Cells stably expressing H1.2/H1.0-Halo or H2B-Halo were cultured on poly-L-lysine coated glass-based dishes (3970-035, Iwaki). HaloTag-tagged molecules were fluorescently labeled with 30-50 pM HaloTag TMR ligand (G8251, Promega), 40 nM PA-JF646 HaloTag (for dual-color imaging) or 20 nM PA-JF549 HaloTag (for Mph imaging) (provided by the Lavis lab at the Janelia Research Campus, VA, USA) for 20 min, washed three times with HBSS buffer (H1387; Sigma-Aldrich), and then imaged in live cells in phenol red free DMEM with 10% FBS.

### Single molecule imaging microscopy

Single molecules were observed using an inverted Nikon Eclipse Ti microscope with a 100 mW Sapphire 561-nm laser (Sapphire-561-100CW CDRH, Coherent) and the scientific complementary metal-oxide semiconductor (sCMOS) ORCA-Fusion BT camera (C15440-20UP, Hamamatsu Photonics). Fluorescently labeled molecules in living cells were excited by the 488-nm, 561-nm, or 640-nm lasers through an objective lens (100× Apo TIRF, NA 1.49; Nikon) and their emissions were detected. An oblique illumination with a TIRF unit (TI-TIRF-E, Nikon) was used to excite labeled molecules within a thin area in the cell nucleus and reduce the background noise. Hundreds of sequential image frames were acquired using MetaMorph (Molecular Devices) or NIS Elements (Nikon Solutions) at 50 ms per frame under continuous illumination. To maintain cell culture conditions (37°C, 5% CO_2_, and humidity) under the microscope, a live cell chamber with a digital gas mixer and a warming box (Tokai Hit) were used.

### Single-molecule tracking and analysis

Sequential microscopy images were converted to 16-bit grayscale and the background signals were subtracted with the rolling ball background subtraction (radius, 50 pixels). The nuclear regions in the images were manually extracted. Single molecules detection and tracking were performed using the Fiji package (TrackMate (*81*)) with the following parameters (Detector: LoGdetector, Object diameter: 7-pixel, Quality threshold: 2, Sub-pixel localization, Initial thresholding: 7-60 (variable depending on the experiment), Tracker: LAP Tracker, Linking distance: 10 pixel or 2pixel). For single-nucleosome movement analysis, the displacement distribution and the MSD of the fluorescent dots were calculated on the basis of their trajectory using a Python program. The originally calculated MSD was in 2D. To obtain the 3D value, the 2D value was multiplied by 1.5 (4 to 6 Dt). Graphs of the obtained single nucleosome MSD between various conditions were performed using R or Python. To ascertain the position determination accuracy of the molecules labeled with TMR in FA-fixed cells, distributions of nucleosome displacements from the centroid of the trajectory in the x- and y-planes in the 50-ms interval were fitted to Gaussian functions. The mean of SD was used for position determination accuracy.

### Motion classification by vbSPT (variational Bayes Single Particle Tracking)

Trajectories longer than 4 frames were used as input to the vbSPT algorithm (*82*) in MATLAB, by imposing a maximum number of states equal to 3 or 2. The algorithm provided the transition rates between these states. For accurate estimation, the diffusion coefficient (*D*) for each state was calculated from the 10 ms/frame data. Also, based on the predicted model, steps in trajectories were annotated into each state. The annotation was used for splitting trajectories that only contained each state.

### Dual color imaging (PALM + single molecule imaging)

Single molecules were observed using an inverted Nikon Eclipse Ti microscope with an ILE laser unit (ANDOR) and the sCMOS ORCA-Fusion BT camera. Fluorescent molecules in living cells were excited by the 561-nm or 640-nm laser through an objective lens (100× Apo TIRF, NA 1.49; Nikon) and detected. A weak (0.1%) 405-nm laser was also used for photoactivation of PAmCherry and PA-JF646. An oblique illumination system with a TIRF unit (TI2-LA-TIRF-E, Nikon) was used to excite labeled molecules within a limited thin area in the cell nucleus and reduce the background noise. Sequential 1000 images were acquired using NIS-Elements (Nikon solutions) at a frame rate of 50 ms under continuous illumination. To maintain cell culture conditions (37°C, 5% CO_2_, and humidity) under the microscope, a live cell chamber with a digital gas mixer and a warming box (Tokai Hit) were used. Beam splitter W-VIEW GEMINI (Hamamatsu Photonics) was used with filters. Chromatic shifts of channels were aligned manually with quad-fluorescent beads.

### Dual color imaging (single molecule imaging + mitotic chromosome)

H1.2-Halo was labeled with PA-JF549 HaloTag. To stain mitotic chromosomes in live cells, 100 nM SiR-Hoechst (SiR-DNA, #CY-SC007; Spirochrome) was added to the medium. Fluorescent molecules in living cells were excited by the 561-nm or 640-nm laser. Beam splitter W-VIEW GEMINI (Hamamatsu Photonics) was used with filters. Cells were asynchronous.

### Co-localization analysis of chromatin domain and H1

PALM images were constructed using the Fiji package ThunderSTORM (*107*) with 2x magnification based on sequential images of PAmCherry, resulting in 32.5 nm x 32.5 nm for a pixel and 740 px x 740 px image. Middle of the nucleus: 220 px x 220 px was cropped for this analysis to avoid the nuclear periphery. All pixels were classified into four classes by quad percentile based on the intensity distribution. From classified trajectories by vbSPT, each middle point of a step belonged to a pixel, so the total number of steps could be calculated for each class. Enrichment for each class (number of steps with the state included in the class/number of all steps with the state) was calculated for single-cell data, then data from more than five cells was averaged for results. If steps were distributed randomly, each class should contain 25% of steps, so that enrichment is colored with 0.25 as a border. A PALM image and trajectories were visualized by a Python script.

### PALM imaging for fixed cells

Cells stably expressing H2B-Halo were cultured on poly-L-lysine coated glass-based dishes (3970-035, Iwaki). HaloTag-tagged molecules were fluorescently labeled with 200 nM HaloTag PA-JF646 ligand (provided by the Lavis lab, Janelia Research Campus) for 20 min. Cells were washed three times with HBSS buffer (H1387; Sigma-Aldrich) and then incubated with phenol free McCoy’s 5A medium (49431-32, KANTO CHEMICAL) with 10% FBS and 0.01% DMSO or 10% FBS and 1 µM 5-Ph IAA for 3 hours. Cells were fixed with 3.7% formaldehyde (FA) (064-00406, Wako) in Opti-MEM (31985062, Gibco) at room temperature for 15 min, washed three times with 1X PBS, and observed in phenol red free McCoy’s 5A medium with 10% FBS. Here, PA-JF646 was illuminated with a stronger (640-nm) and a weaker (405-nm) laser. H1.2-mClover signal was also detected using a 488-nm laser.

### Clustering analyses of nucleosomes in PALM images

The methods for clustering analyses of nucleosomes in PALM images were described previously (*8*). Ripley’s *K* function is given by

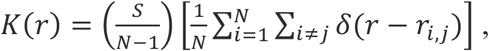

where (*N* – 1)/*S* is the average particle density of area *S*, and *N* is the total number of particles contained in the area. The delta function is given by

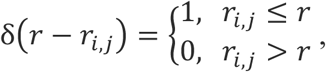

where *r_i_*_,*j*_ is the distance between *r_i_* and *r_j_*.

The *L* function is given by

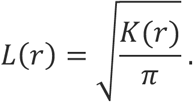

The area *S* of the total nuclear region was estimated using the Fiji plugin Trainable Weka Segmentation, and the area of the whole region was measured by Analyze Particles (ImageJ).

### Expression of wild-type H1.2 and mutated H1.2 in HCT116 cells

Either wildtype H1.2-HaloTag (H1.2-WT) or mutated H1.2-HaloTag (H1.2-Khalf) was transiently expressed using the Tet-On inducible system. Inducible vectors of H1.2-WT or H1.2-Khalf were transfected into HCT116 with H1.2-mAC using the Neon electroporation transfection system and seeded on a glass bottom dish. Two days after transfection, cells were treated with 1 µM 5-Ph IAA and doxycycline (1 µg/mL) for 6 hours to deplete endogenous H1.2 and ectopic expression of H1.2-WT or H1.2-Khalf was induced. Then, HaloTag-tagged H1.2-WT or H1.2-Khalf were fluorescently labeled with 10 pM JFX650 HaloTag ligand (HT1070, Promega) and 100 nM HaloTag TMR ligand for 20 min. The cells were washed three times with HBSS buffer (H1387; Sigma-Aldrich), then observed in phenol red free McCoy’s 5A medium with 10% FBS and 1 µM 5-Ph IAA and doxycycline (1 µg/mL). Red fluorescence from TMR was used for searching cells that expressed H1.2-WT or H1.2-Khalf, and far-red fluorescence from JFX650 was used for single molecule tracking.

### The Richardson and Lucy (RL) algorithm

We utilized the Richardson and Lucy (RL) algorithm to derive a smooth distribution curve from the sparse and noisy histogram of the single-cell molecule trajectory data (*84*). The single-cell distributions of the two-dimensional mean squared displacement (MSD), denoted by *M*, are shown in Figs. 3B and S5E. These distributions were obtained by numerically iterating the RL calculation. To start, we sampled the two-dimensional position 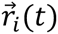 of the *i*th molecule of the *α*th cell (*i* ∈ *α*) at time *t*. We then calculated the self-part of the van Hove correlation function,

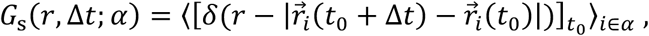

where *δ*(⋯) represents Dirac’s delta function, 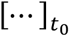 denotes the average over different values of *t*_0_, and 〈⋯ 〉_i∈*α*_ indicates the average over the molecules *i* belonging to the cell *α*. The iteration started from an initial estimate for *P*(*M*, Δ*t*; *α*) as *P*^0^(*M*, Δ*t*; *α*) = exp(− *M*⁄*M*_0_) /*M*_0_, where the convergent results were not sensitive to the choice of the *M*_0_ value. The RL calculation was iterated using

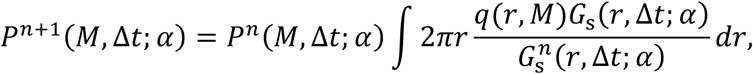

under constraints *P^n^*^+1^(*M*; *α*) ≥ 0 and ∫ *dM P^n^*^+1^(*M*; *α*) = 1. Here, *q*(*r*, *M*) = exp(− *r*^1^⁄*M*) /(*πM*) and 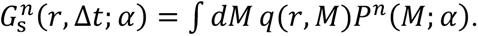 The process was continued until the difference between *P^n^*^+1^(*M*; *α*) and *P^n^*(*M*; *α*) became sufficiently small, resulting in the convergent distribution *P*(*M*, Δ*t*; *α*) = *P^n^*^+1^(*M*, Δ*t*; *α*). While the quantitative features of *P*(*M*, Δ*t*; *α*) vary from cell to cell, the qualitative features̶including the number of peaks and the rough difference among peak positions̶remain consistent across different cells. Therefore, distributions *P*(*M*, Δ*t*; *α*) depicted for an example cell *α* in Figs. 3B and S5E reflect the essential characteristics common to the *n* = 40 cells examined.

### The mean-field theory of H1 and nucleosomes

The multiscale modeling of interactions between H1 molecules and nucleosomes (Figs. 1 and 5) demonstrated that H1 can interact with multiple nucleosomes in a short period, leading to weak multivalent interactions. We anticipate that these interactions are associated with both a decrease in energy and an increase in entropy. To investigate whether these multivalent interactions and the resulting entropy gain stabilize the dynamic H1 molecules, we examined a mean-field model similar to the lattice polymer model of Flory and Huggins (*93*) (Fig. S9A). In this model, we assume that chromatin chains, composed of *K* nucleosomes, are placed on the lattice with *N*_0_, grid points. Each grid point can either be occupied by a nucleosome or be left unoccupied, resulting in a total of *N* nucleosomes; therefore, the nucleosome density is given by *N*/*N*_0_. H1 is modeled to jump over the lattice sites with a chemical potential *μ* when it is not bound to a nucleosome. Each nucleosome can either bind one H1 molecule or remain unbound. When H1 binds at the most stable dyad position of a nucleosome, H1 is stabilized by an energy of −(*ε* + *δ*) with *ε* > 0 and *δ* > 0. If H1 dynamically binds near the dyad, it has a higher binding energy of −*ε* < 0. We assume that such a dynamically bound H1 can interact with nucleosomes at neighboring lattice sites, with an interaction energy of −*J* < 0. Then, the mean-field energy can be expressed as

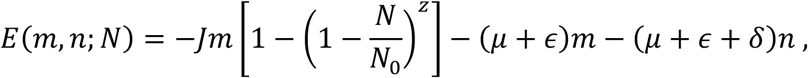

where *m* represents the number of H1 molecules with dynamic binding, *n* is the number of H1 molecules with stable binding, and *z* is the number of neighboring lattice sites. The first term on the right-hand side of this equation accounts for the multivalent interactions between H1 and nucleosomes, representing the nonlinear dependence of energy on *N*⁄*N*_0_. Entropy is derived from the number of states, estimated as

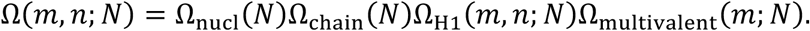

Here, 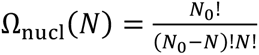 shows the effects of nucleosome motion, 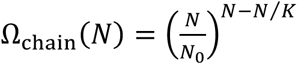 accounts for the connection in the chromatin chain, 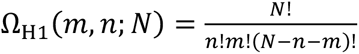 shows the distribution of H1 bound to nucleosomes, and 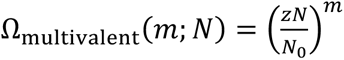
represents the fluctuation in multivalent interactions between the mobile H1 and neighbor nucleosomes. Entropy *S* = *k_B_*logΩ with *k_B_* being the Boltzmann factor, is

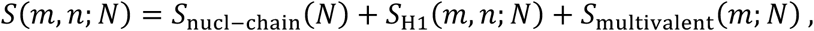

with *S*_nucl-chain_(*N*) = *k_B_*log/Ω_nucl_(*N*)Ω_chain_(*N*)0, *S*_H1_(*m*, *n*; *N*) = *k_B_*logΩ_H1_(*m*, *n*; *N*), and *S*_multivalent_(*m*; *N*) = *k_B_*logΩ_multivalent_(*m*; *N*). Neglecting the higher order terms of 1⁄*N*_0_, each componet of entropy is

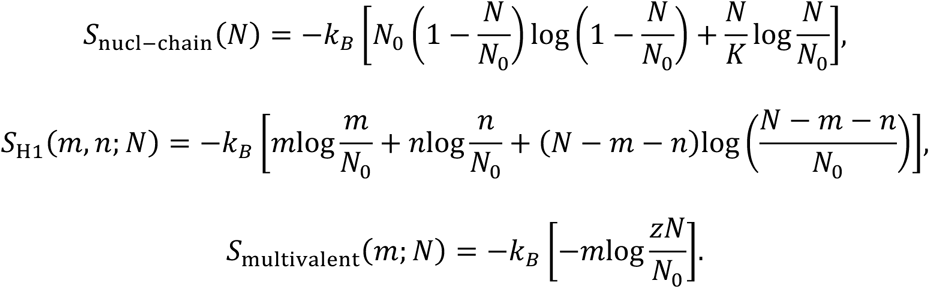

Here, *S*_multivalent_(*m*; *N*) represents the entropy effect of multivalent interactions of dynamic H1. Then, the free energy per lattice site at temperature *T* is given by

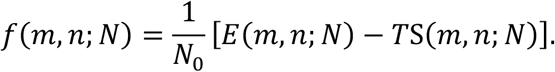

In Fig. S9B, *f*(*m*, *n*; *N*) is plotted on a two-dimensional plane representing the ratio of the H1 on nucleosome, (*n* + *m*)⁄*N*, and the ratio of H1 with dynamic binding, *m*⁄(*n* + *m*). To represent the stable binding of H1 at the dyad position of nucleosomes, we assume *δ* > *ε* ≈ *μ* ≈ *J* > 0.

The parameters used for the results in Fig. S9 were chosen as *K* = 1000, *z* = 6, *δ*⁄*k_B_T* = 1.5, and *ε*⁄*k_B_T* = *μ*⁄*k_B_T* = *J*⁄*k_B_T* = 1.

## Author contributions

Project design: MAS, SI, RC-G, KM

Generated H1.2/H1.0-Halo expressing cells: SI

Single-nucleosome imaging, fluorescence imaging, and analyses: MAS, SI

Computational modeling: JH, CP, SF, RC-G

Biochemical experiments and creation of HCT116 AID cells with help of SI: ST

Analysis development: SSA and MS

Writing – original draft: MAS, JH, RC-G, KM

Writing – review & editing: all authors

## Acknowledgments

We are grateful to Ms. K. Sato for establishing H1.2/H1.0-Halo expressing cell lines and the initial analysis, Dr. K. M. Marshall, Ms. K. Nakazato, and Mr. Y. Nagata for critical reading and editing of this manuscript, Dr. C. A. Davey and Dr. J. C. Hansen for the collaboration on biochemical analysis of H1-nucleosome array condensates, and Dr. M. T. Kanemaki for providing HCT116 cells. We thank Dr. K. Hibino, Dr. S. Hirose, Dr. K. Kurokawa, Dr. Y. Shimamoto, Dr. T. Torisawa, and Maeshima laboratory members for helpful discussions and support. This work was supported by the following funding sources:

Japan Society for the Promotion of Science and MEXT KAKENHI grant JP23K17398 (S.I. and KM) Japan Society for the Promotion of Science and MEXT KAKENHI grant JP24H00061 (KM and MS)

Japan Society for the Promotion of Science KAKENHI grant JP22H00406 (MS)

Japan Society for the Promotion of Science and MEXT KAKENHI grant JP21H02535 (SI)

Japan Society for the Promotion of Science and MEXT KAKENHI grant JP22H05606 (SI)

Takeda Science Foundation (KM)

UK Government’s Guarantee scheme EP/Z002028/1 following funding from the European Research Council (Consolidator Grant)(RC-G)

UK High-End Computing Consortium for Biomolecular Simulation EP/R029407/1 (RC-G) Herchel Smith Postdoctoral Fellowship (JH)

UK Research and Innovation (UKRI) Postdoctoral Fellowships Guarantee scheme EP/X02332X/1 (JH)

Japan Society for the Promotion of Science Fellow JP24KJ1161 (MAS)

National Institute of Genetics 2023 NIG-JOINT 4I2023 (RC-G and KM)

## Competing interests

The authors declare no competing interests.

## Data and material availability

All data are available in the main text or the supplementary materials.

## Supplementary Materials

**Fig. S1.**
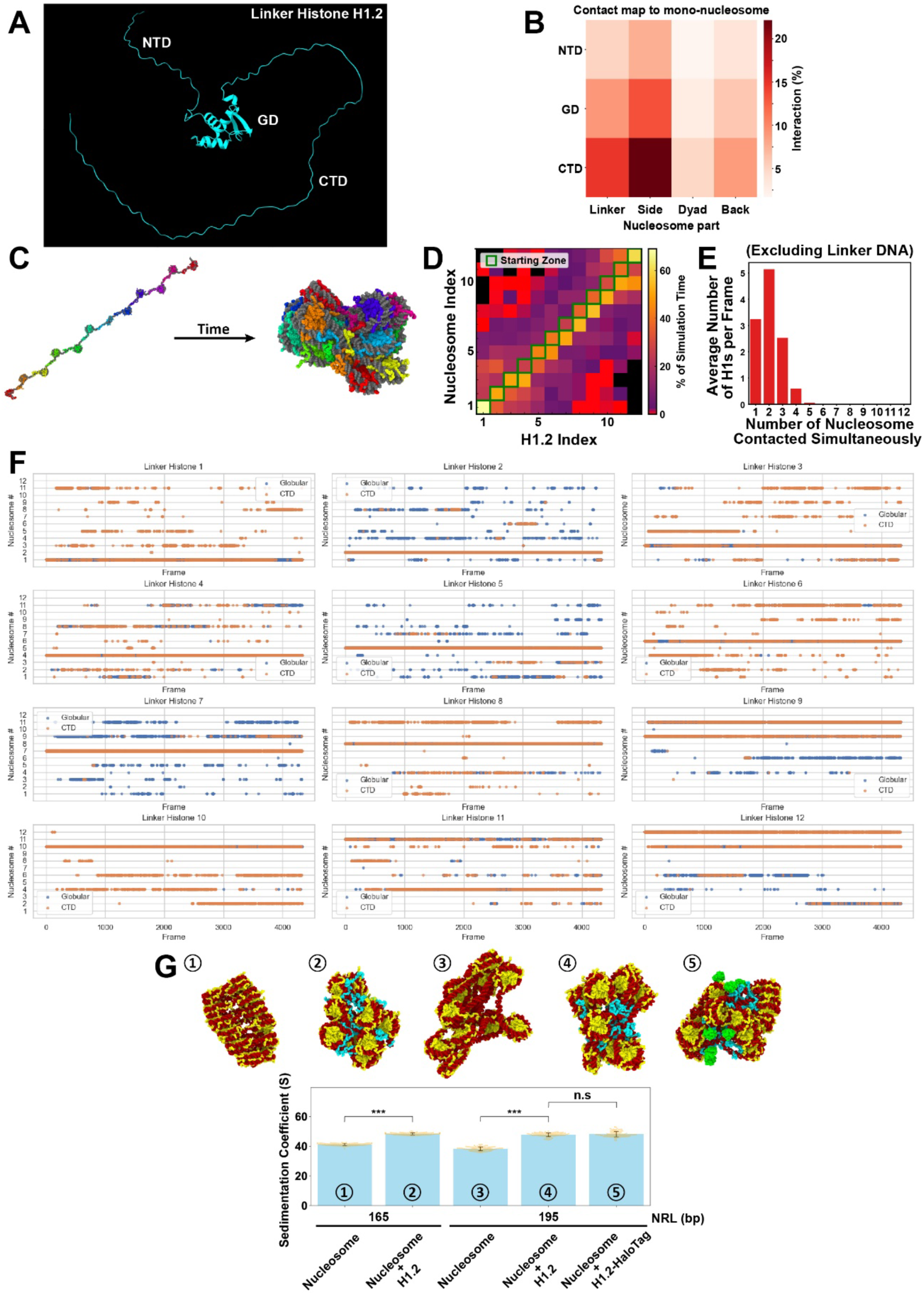
Models of a linker histone H1 and a 12-mer nucleosome array. **(A)** Structure of human linker histone H1.2 predicted by AlphaFold3: unstructured N-terminal domain (NTD), globular domain (GD), and a long, intrinsically disordered C-terminal domain (CTD). **(B)** Region-specific contact map between a H1.2 molecule and a mono-nucleosome. H1.2 (y-axis): NTD, GD, CTD regions; Nucleosome (x-axis): Linker, Side, Dyad, and Back regions. **(C)** Initial configuration (left) and final configuration (right) of a 12-mer nucleosome array (195 bp NRL) with H1.2 (Nucleosome:H1 = 1:1). Originally, each H1.2 was attached to one nucleosome on a linear array, then H1.2 translocated to other nucleosomes over time, twisting and condensing the array. **(D)** Contact map between twelve nucleosomes and twelve H1.2 on the array (195 bp NRL). Starting zones on the diagonal (square edge in green) indicate where H1.2 is located at the beginning of the simulation. Note that H1.2 often left the original nucleosome and translocated among several nucleosomes. **(E)** Histogram of the number of nucleosomes contacted simultaneously by an H1.2. Only core nucleosomal DNA was taken into account. **(F)** Most-contacted nucleosome (excluding linker DNA) in an unbiased MD simulation of a 12-mer nucleosome array with 195 bp NRL. For each linker histone, the nucleosome with the highest average number of contacts per frame is shown separately for the GD and CTD. **(G)** Examples of nucleosome array configurations and their corresponding sedimentation coefficients (S) (± standard deviation, SD) with different conditions. With H1.2, the nucleosome arrays show a significantly higher S, indicating greater compaction. Similar trends were observed for different NRLs or with a H1.2-HaloTag (green sphere). The Wilcoxon rank-sum test was used to determine P values and corrected by the Benjamini-Hochberg method. ***P < 0.0001 for nucleosomes with 165 bp NRL versus nucleosomes with 165 bp NRL + H1.2 (P = 4.2 × 10^−34^), nucleosomes with 195 bp NRL versus nucleosomes with 195 bp NRL + H1.2 (P = 4.2 × 10^−34^). Not significant (N.S.) for nucleosomes with 195 bp NRL + H1.2 versus nucleosomes with 195 bp NRL + H1.2-Halo (P = 0.89).

**Fig. S2.**
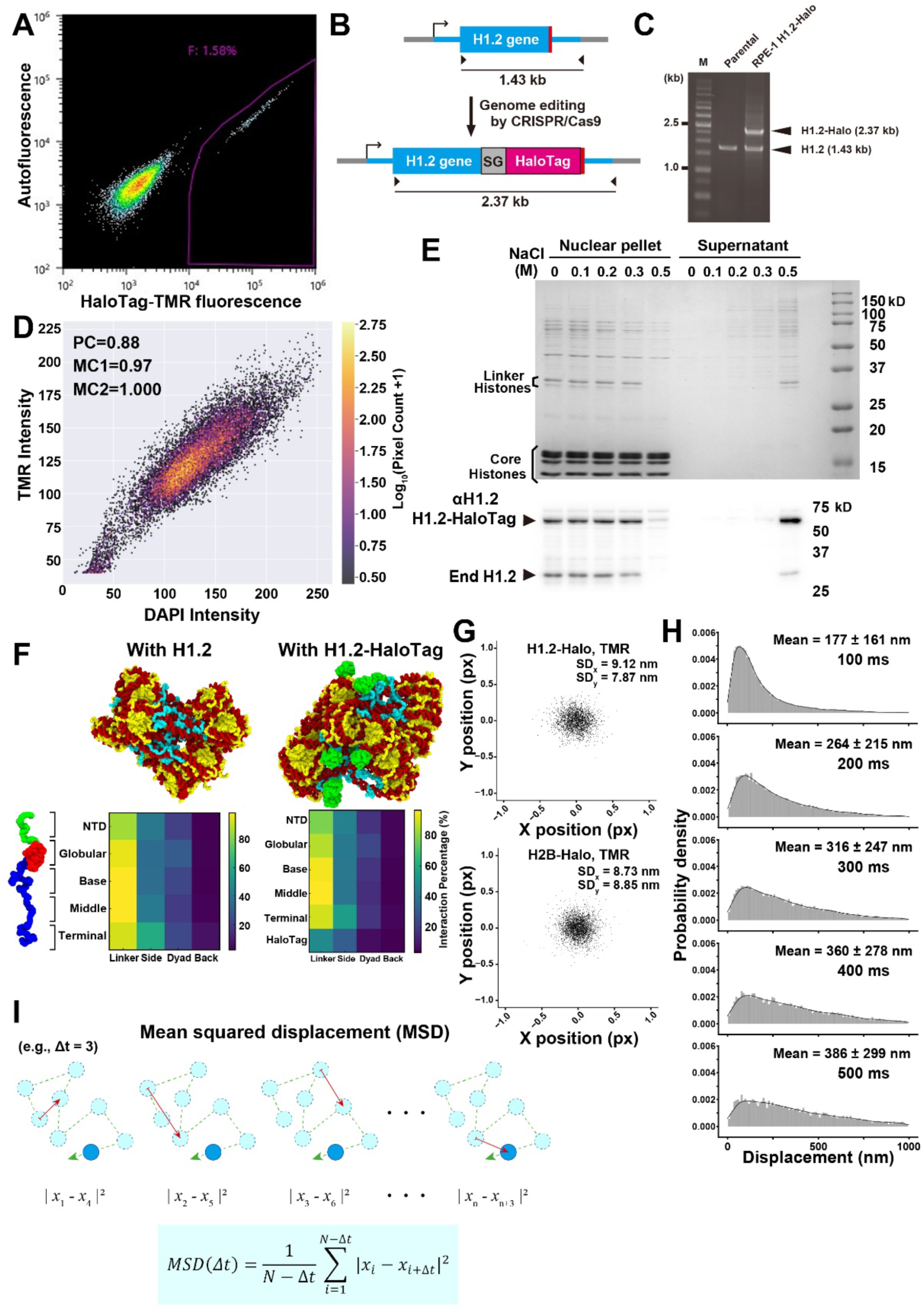
Generation of RPE-1 cells expressing the H1.2-HaloTag and characterization of H1.2-Halo behavior. **(A)** Intensity scatter plot of FACS for sorted RPE-1 cells expressing H1.2-HaloTag (H1.2-Halo). The gate used to collect Halo-TMR positive cells is shown with a pink line. **(B)** Scheme for the original and HaloTag-inserted H1.2 gene loci and expected fragments amplified by PCR with the indicated primer set. **(C)** Validation for proper mono-allelic insertion of the HaloTag in RPE-1 genomic DNA by PCR: Parental (left) and H1.2-Halo (right) RPE-1 cells. HaloTag was inserted into the heterozygous H1.2 gene loci. **(D)** Pixel intensity correlation of DAPI and TMR staining in Fig. 2B. PC, Pearson correlation coefficient; MC1/MC2, Manders correlation coefficients. **(E)** Stepwise-salt washing of nuclei isolated from the H1.2-Halo expressing RPE-1 cells. The nuclei were washed with buffers containing increasing concentrations of NaCl. The resultant nuclear pellets (left) and supernatants (right) were analyzed by SDS-PAGE, and subsequently stained with Coomassie brilliant blue (top) or immunoblotted with anti-H1.2 (bottom). Positions of core histones and linker histone H1 are indicated on the Coomassie brilliant blue stained gel. Note that H1.2 and H1.2-Halo dissociate from chromatin with 0.5 M NaCl and were detected in the supernatant fraction, suggesting that H1.2-Halo interacts with chromatin like endogenous H1.2. **(F)** Representative configuration and region-specific contact map of 12-mer nucleosome array with H1.2 or H1.2-Halo. Note that both conditions showed compact, irregular clusters and similar region-specific contact maps. **(G)** Scatter plot of H2B-Halo-TMR (n = 214) and H1.2-Halo-TMR (n = 155) dots in formaldehyde (FA)-fixed cell to ascertain the position determination accuracy. Standard deviations on the x-axis and y-axis are shown as SD_x_ and SD_y_. **(H)** Displacement distribution histograms (n = 40 cells) for 100, 200, 300, 400, and 500 ms. Means ± SD of displacement are at the top of each. **(I)** Calculation of mean squared displacement (MSD).

**Fig. S3.**
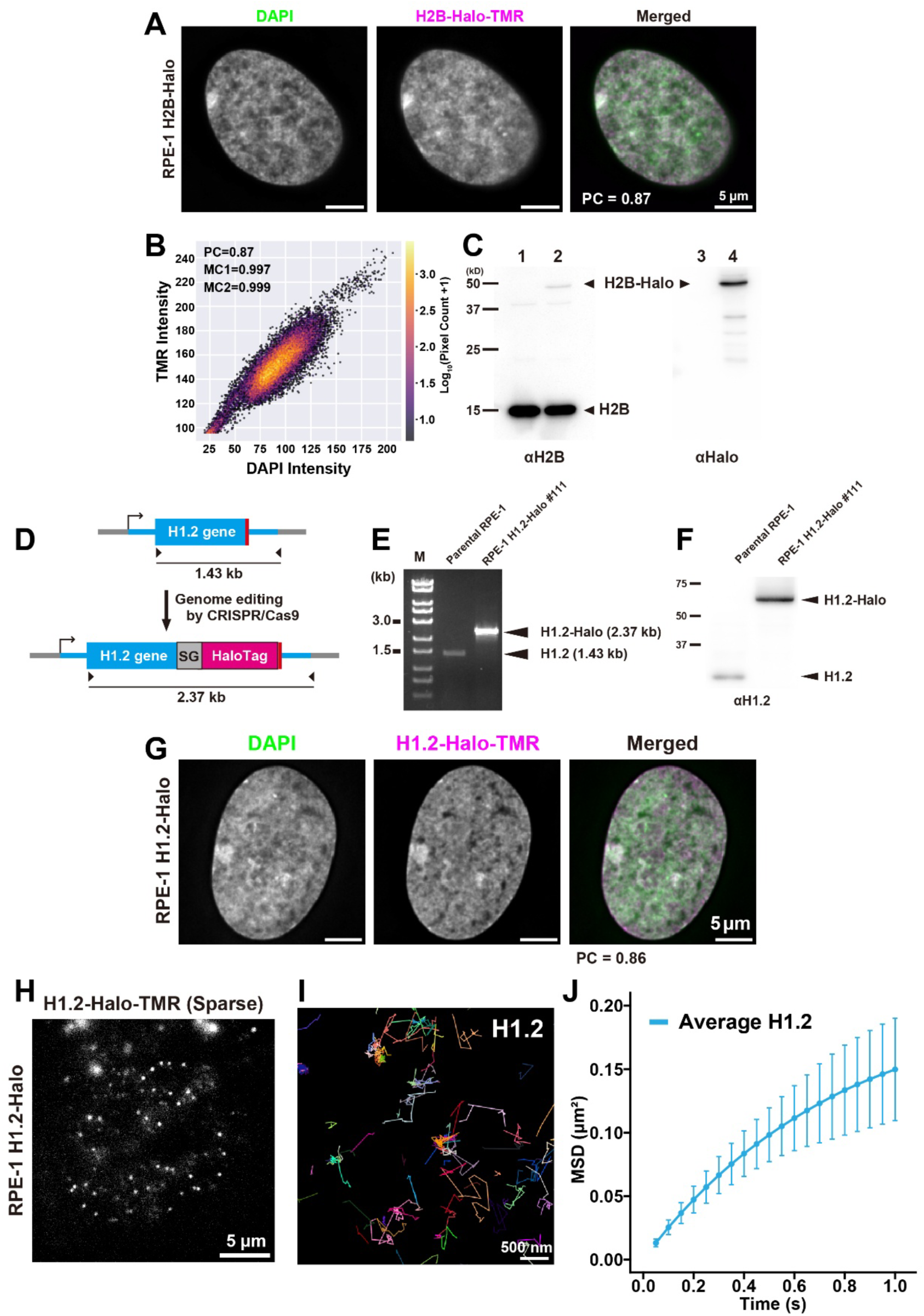
Generation of RPE-1 cells ectopically expressing H2B-Halo and of RPE-1 cells expressing H1.2-Halo via bi-allelic HaloTag insertion. **(A)** Representative images of RPE-1 cells ectopically expressing H2B-Halo. The localization of H2B-Halo-TMR is similar to that of DAPI. **(B)** Pixel intensity correlation of the DAPI and TMR images in Fig. S3A. PC, Pearson correlation coefficient; MC1/MC2, Manders correlation coefficient. **(C)** Western blotting of parental RPE-1 (lanes 1, 3) and RPE-1 expressing H2B-Halo (lanes 2, 4) using anti-H2B antibody (lanes 1, 2) and anti-HaloTag antibody (lanes 3, 4). **(D)** Scheme for the parental and HaloTag-inserted H1.2 gene loci and expected fragments amplified by PCR with the indicated primer set. **(E)** Validation for bi-allelic insertion of the HaloTag in RPE-1 genomic DNA by PCR: Parental (left) and H1.2-Halo (right) RPE-1 cells. **(F)** Western blotting of parental RPE-1 (left) and RPE-1 expressing H1.2-Halo (right) using an anti-H1.2 antibody. Note that all H1.2 is tagged by HaloTag. **(G)** Representative images of RPE-1 cells expressing H1.2-HaloTag via bi-allelic HaloTag insertion. The localization of H1.2-Halo-TMR is similar to that of DAPI. **(H)** A representative single-molecule image of H1.2-Halo-TMR in the generated RPE-1 cells. Each white dot represents a single molecule of H1.2-Halo. **(I)** Representative trajectories of single nucleosomes labeled with H1.2-Halo-TMR (right) acquired at 50 ms/frame. **(J)** Mean squared displacement (MSD) plots (± SD among cells) of single H1.2 molecules (blue) in living RPE-1 cells in a tracking time range from 0.05 to 1 s (n = 30 cells).

**Fig. S4.**
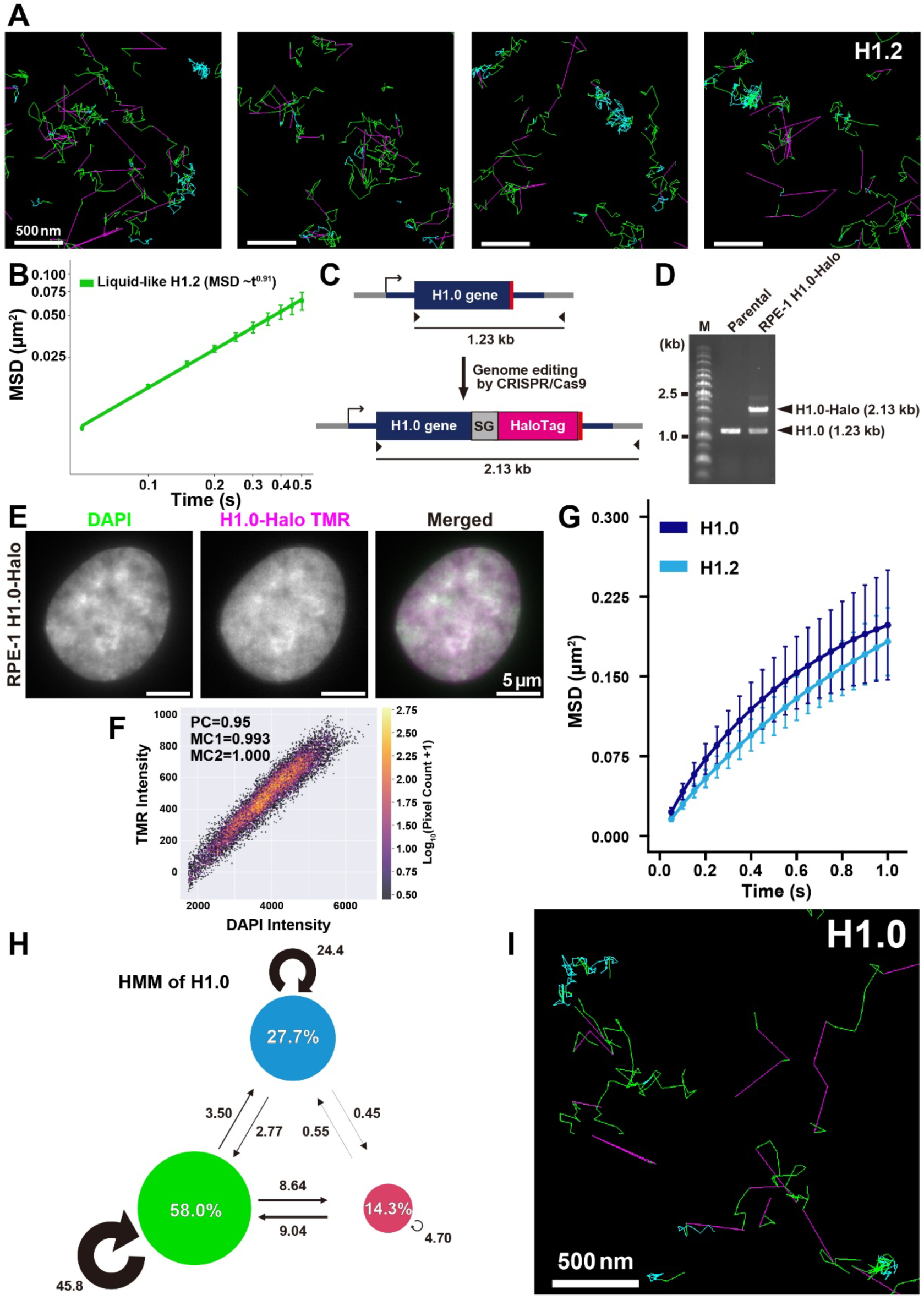
Further trajectory analysis of H1.2, creation of RPE-1 cells expressing H1.0-Halo, and single-H1.0 imaging. **(A)** Examples of classified trajectories of H1.2. **(B)** The log-log plot of MSDs from the plot of “liquid-like H1”. The plot was fitted linearly. The calculated anomalous exponent is also shown. **(C)** Schematic of CRISPR/Cas9-mediated genome editing for inserting the HaloTag at the C-terminus of the H1.0 gene locus. SG, linker (GGGGS x2). Expected PCR fragment lengths for original and HaloTag-inserted H1.0 gene loci are shown. **(D)** Validation for proper insertion of HaloTag in RPE-1 genomic DNA by PCR: Parental (left) and H1.0-Halo (right) RPE-1 cells. HaloTag was inserted into the heterozygous H1.0 gene locus. **(E)** Representative images of FA-fixed RPE-1 cells expressing H1.0-Halo. The localization of H1.0-Halo-TMR is similar to that of DAPI. **(F)** Pixel intensity correlation of DAPI and TMR images in Fig. S4E. PC, Pearson correlation coefficient; MC1/MC2, Manders correlation coefficient. **(G)** MSD plots (± SD among cells) of single H1.0 and H1.2 in living RPE-1 cells over a tracking time range from 0.05 to 1 s (n = 40 cells for each)(H1.2 data was reproduced from Fig. 2H). **(H-I)** vbSPT classification of H1.0 trajectories (n = 40 cells).

**Fig. S5.**
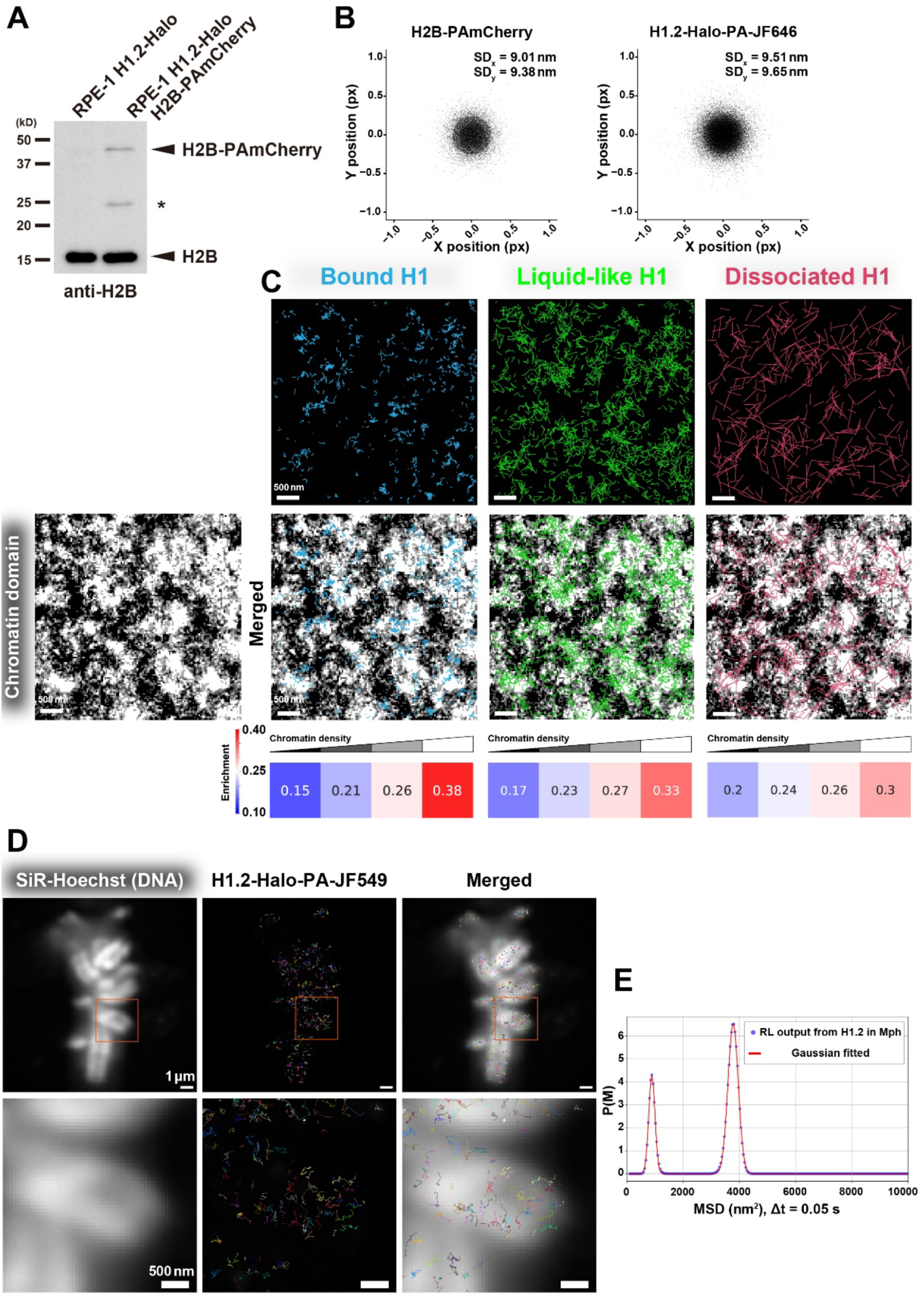
Dual color imaging of H2B-PAmCherry and H1.2-Halo-PA-JF646, and H1.2 movements in metaphase chromosomes. **(A)** Western blotting of parental RPE-1 and RPE-1 H1.2-Halo cells expressing H2B-PAmCherry using an anti-H2B antibody. **(B)** Scatter plots of H2B-PAmCherry (n = 8927) and H1.2-Halo-PA-JF646 (n = 5236) dots in a FA-fixed cell to ascertain the position determination accuracy. Standard deviations on the x-axis and y-axis are shown as SD_x_ and SD_y_. **(C)** Overlap of the PALM image of nucleosomes and trajectories of each state. The distribution of trajectories in each class is shown as enrichment values and colors. A value of 0.25 represents a random distribution of trajectories. Bound H1 and liquid-like H1 show prominent enrichment in chromatin-dense regions. **(D)** Another example of the overlap of SiR-Hoechst (DNA) signal and trajectories of H1 in metaphase chromosomes. Individual trajectories are randomly colored. Bottom panels are enlarged images from squared regions in top panels. Note that H1 shows diffusive movement within mitotic chromosomes. **(E)** Two prominent peaks were obtained from the MSD data of H1.2 in metaphase chromosomes using the RL algorithm (*84*). The outputs from the RL algorithm (blue dots) and the fitted Gaussian mixture (red line) are shown.

**Fig. S6.**
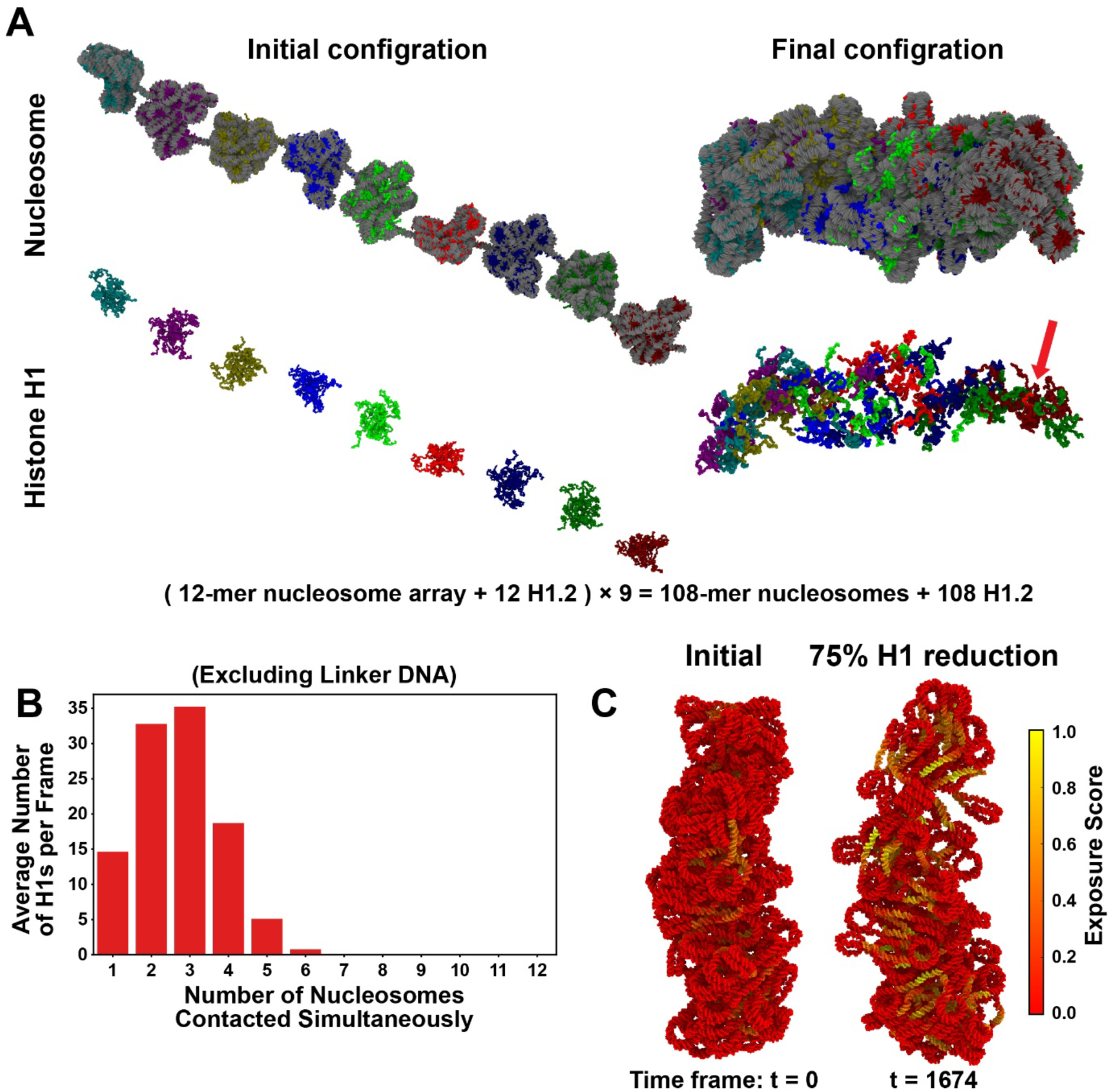
H1.2 behaves like a liquid-like glue within 108 nucleosomes. (A, left) Nine of the 12-mer nucleosome arrays (195 bp NRL) with H1.2 (one H1 per nucleosome) are connected into a fiber at the initial time point. (right) The final configuration shows an irregular cluster of 108 nucleosomes with 108 H1.2 molecules. Nucleosomes and their initially associated H1.2 are color-matched. The bottom panels display only H1. The red arrow indicates a red H1 diffuses away from its original position. Green H1, like red H1, is also dispersed at the final irregular cluster configuration, suggesting the liquid-like behavior of H1 within a dense chromatin domain. **(B)** Histogram of the number of nucleosomes contacted simultaneously by an H1.2 during the 108-nucleosome cluster simulation. Only core nucleosomal DNA was taken into account. **(C)** When 75% of H1.2 was removed from the 108-nucleosome cluster, the cluster became decondensed and DNA was more exposed as shown with exposure score (left, initial configuration; right, final configuration after H1 reduction).

**Fig. S7.**
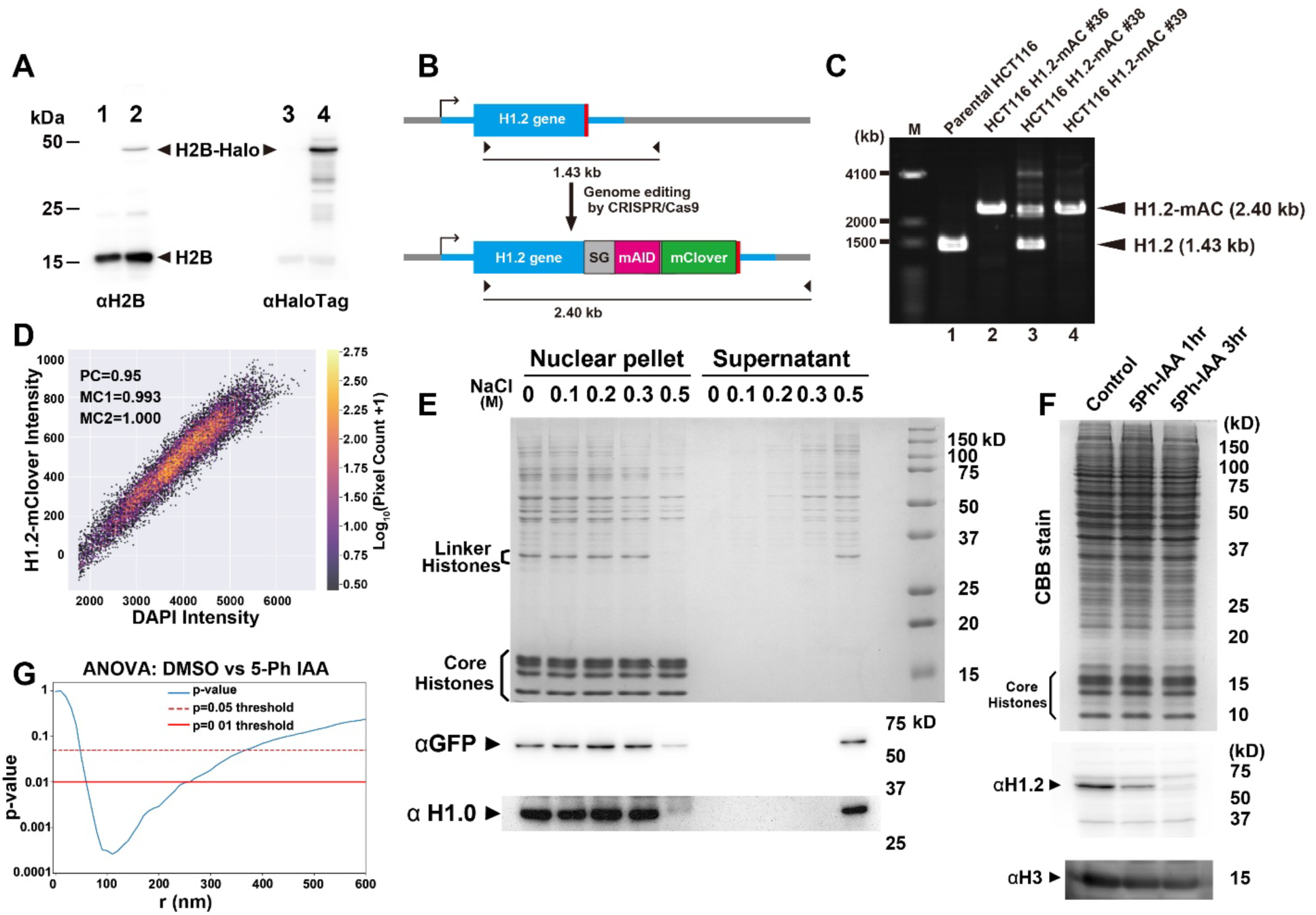
Characterization of HCT116 cells expressing H1.2-mAC and H2B-Halo. **(A)** Western blotting of HCT116 cells expressing H1.2-mAC (lanes 1 and 3) and HCT116 cells expressing H1.2-mAC and H2B-Halo (lanes 2 and 4) using anti-H2B (lanes 1 and 2) and anti-HaloTag (lanes 3 and 4) antibodies. **(B)** Scheme for original and mAC-inserted H1.2 gene loci and expected fragments amplified by PCR with a primer set (arrow heads). **(C)** Validation for proper insertion of mAC by PCR in Parental (left) and H1.2-Halo (right) HCT116 cells. mAC was inserted into the homozygous H1.2 gene loci. **(D)** Pixel intensity correlation between the DAPI and H1.2-mClover images in Fig. 6C. PC, Pearson correlation coefficient; MC1/MC2, Manders correlation coefficient. **(E)** Stepwise-salt washing of nuclei isolated from HCT116 cells expressing H1.2-mAC. The isolated nuclei were washed with the buffers containing increasing concentrations of NaCl. The resultant nuclear pellets (left) and supernatants (right) were analyzed by SDS-PAGE, and subsequently stained with Coomassie brilliant blue (top) or immunoblotted with anti-GFP, anti-H1.2, or anti-H1 antibodies. Positions of core histones and linker histone H1 are indicated in the Coomassie brilliant blue stain. Note that H1.2 and H1.2-mAC dissociate from chromatin with 0.5 M NaCl and were detected in the supernatant fraction, suggesting that H1.2-mAC interacts with chromatin similar to endogenous H1.2. **(F)** Validation of rapid H1.2 depletion. Western blotting of HCT116 cells expressing H1.2-mAC using an anti-H1.2 antibody. Cells were treated with 0.01% DMSO for 3 hours (left), 1 µM 5-Ph IAA for 1 hour (center), or 1 µM 5-Ph IAA for 3 hours (right). Successful depletion of H1.2 was detected after 3 hours of 5-Ph IAA treatment. **(G)** Plot of p-values calculated for the comparison between the L-function of DMSO-treated cells (control) and 5-Ph IAA-treated cells (H1.2-depleted) in Fig. 6H. A one-way analysis of variance (ANOVA) was used to determine the p-values. Notably, a significant reduction was observed in the range of 50 to 350 nm, corresponding to the size of chromatin domains.

**Fig. S8.**
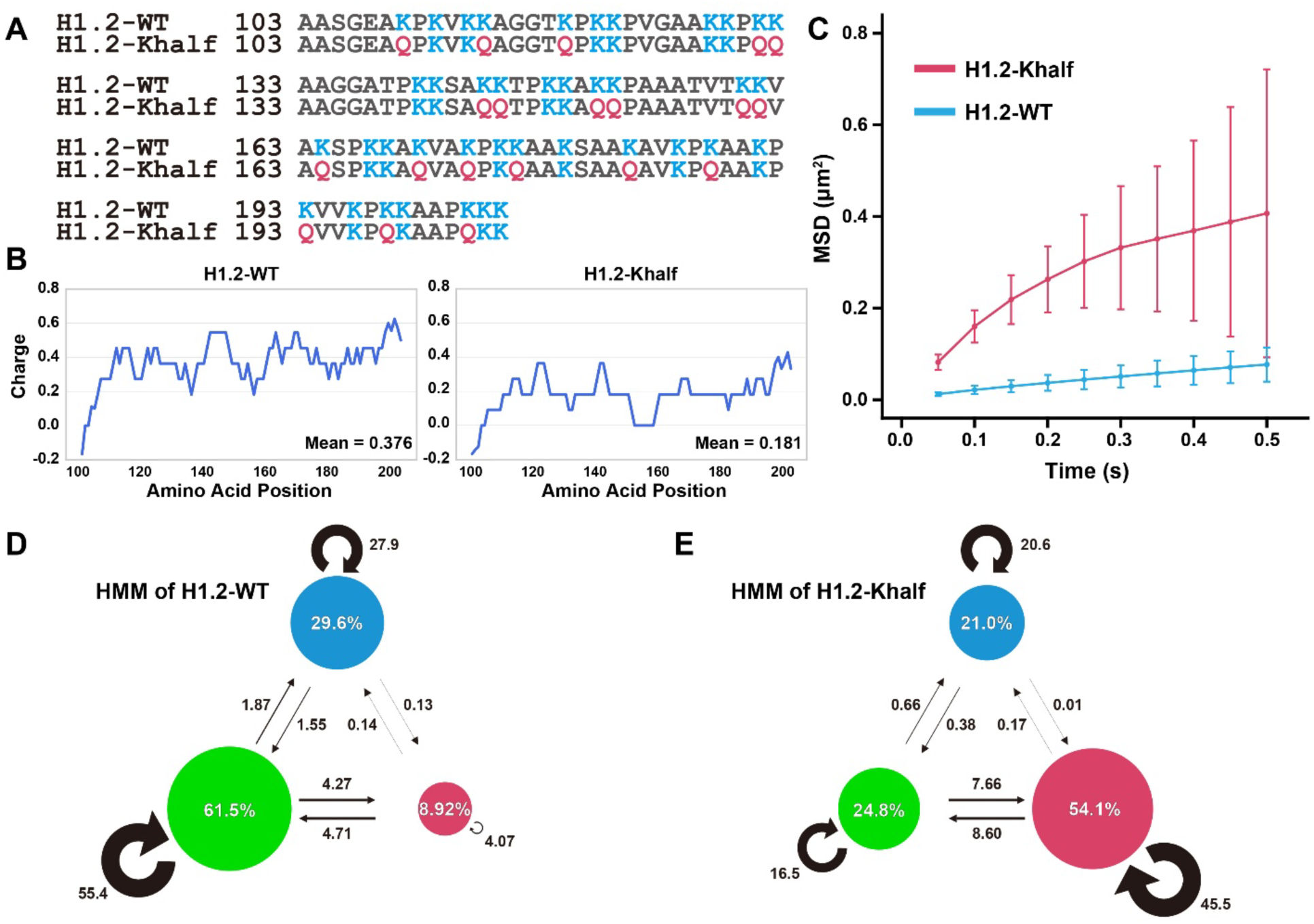
Positive charge on C-terminal IDR is crucial for the liquid-like behavior of H1. **(A)** Comparison of intrinsically disordered CTD amino acid sequences between wild-type H1.2-Halo (H1.2-WT) and mutated H1.2-Halo (H1.2-Khalf). In H1.2-Khalf, half of the lysine residues were replaced with glutamine. **(B)** Charge plots of the intrinsically disordered CTD. Note that H1.2-Khalf had a lower positive charge than the wild type. **(C)** MSD plots (± SD among cells) of H1.2-WT and H1.2-Khalf in living HCT116 cells, tracked over a time range of 0.05 to 0.5 s (n = 20 cells per condition). Both molecules were sparsely labeled with JFX650 HaloTag ligand, and single molecules were tracked. **(D, E)** vbSPT results for H1.2-WT (n = 20 cells) and H1.2-Khalf (n = 20 cells), respectively. WT: *D_Bound_* : 3.26 × 10^-2^ µm^2^/s, *D_Liquid-like_* : 1.05 × 10^-1^ µm^2^/s, *D_Dissociated_* : 1.32 µm^2^/s, Khalf: *D_Bound_* : 3.19 × 10^-2^ µm^2^/s, *D_Liquid-like_* : 1.56 × 10^-1^ µm^2^/s, *D_Dissociated_* : 1.63 µm^2^/s. Notably, the proportion of “liquid-like H1” drastically decreased in H1.2-Khalf compared with H1.2-WT.

**Fig. S9.**
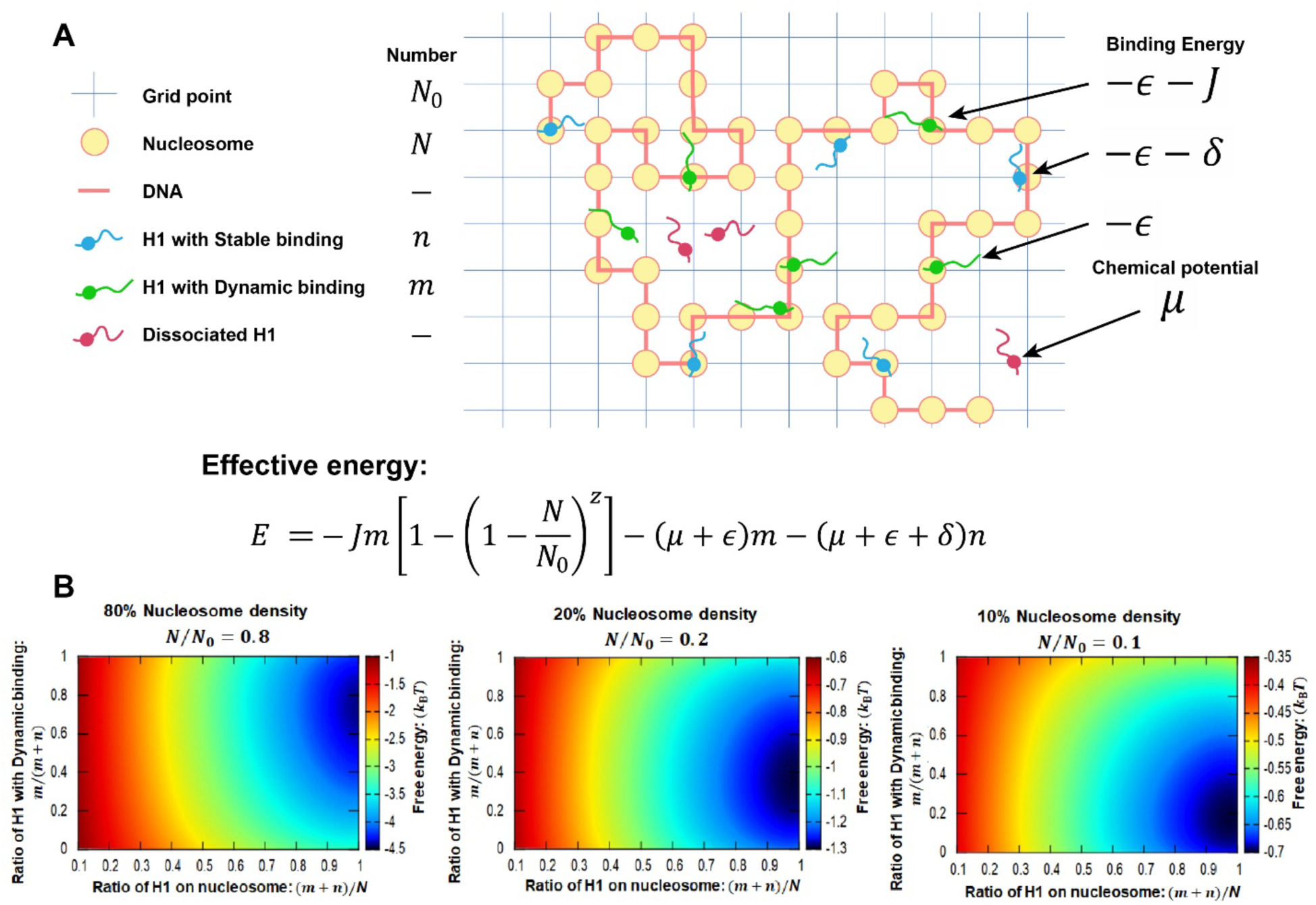
Multivalent interactions and associating entropy increase stabilize the mobile state of interactions between H1 and nucleosomes. **(A)** Schematic representation of the mean-field theory of H1 and nucleosomes (see the Methods for details). *N* nucleosomes are arranged on *N*_0_ grid points. Of the H1 molecules, *n* binds to the most stable sites at the nucleosome dyads, with the intrinsically disordered CTD of each H1 interacting with nearby DNA. These H1 molecules are in a low-energy state of − *ɛ* − *δ* (*ɛ* > 0, *δ* > 0). Other *m* H1 molecules bind near the dyads in higher-energy states of − *ɛ*. If an H1’s intrinsically disordered CTD associates with a nearby nucleosome located within *z* neighboring grid points, the system stabilizes further by − *J* per pair. Dissociated H1 molecules, characterized by chemical potential *μ*, diffuse freely. **(B)** Free energy plot with different nucleosome densities (80%, 20%, 10%). With denser nucleosomes, a higher ratio of H1 with dynamic binding is favored.

## Legends for Movies

**Movie S1.**

MD simulation of a 12-mer nucleosome array with 12 linker histone H1 molecules. The nucleosome repeat length (NRL) is 165 bp. In the presence of H1, the nucleosome array adopts an irregular folding pattern, with H1 dynamically interacting with multiple nucleosomes.

**Movie S2.**

MD simulation of a 12-mer nucleosome array (195 bp NRL) with 12 linker histone H1 molecules. In the presence of H1, the nucleosome array adopts an irregular folding pattern, with H1 dynamically interacting with multiple nucleosomes.

**Movie S3.**

Live-cell imaging of single nucleosomes labeled with TMR in an RPE-1 cell (50 ms per frame). Clear, well-separated dots are observed, each exhibiting single-step photobleaching, indicating that each dot represents a single H2B-Halo-TMR molecule within an individual nucleosome. The scale bar is 5 µm.

**Movie S4.**

Live-cell imaging of single H1.2 molecules labeled with TMR in an RPE-1 cell (50 ms per frame). Clear, well-separated dots are observed, each exhibiting single-step photobleaching after background subtraction (see Fig. 2E). This indicates that each dot represents a single H1.2-Halo-TMR molecule associated with chromatin. The scale bar is 5 µm.

**Movie S5.**

Dual-color live-cell imaging of single H2B-PAmCherry (left) and single H1.2 molecules labeled with PA-JF646 (right) in an RPE-1 cell (50 ms per frame). After PALM reconstruction and tracking, images in Movie S6 were obtained.

**Movie S6.**

Movie of single H1.2 molecules labeled with PA-JF646 in an RPE-1 cell (50 ms per frame). The determined position of each H1.2 molecule was shown as a green particle, while recent trajectories were depicted in cyan. Chromatin domain structures were visualized in magenta using PALM imaging of H2B-PAmCherry. Notably, the “liquid-like” motion of H1 is predominantly observed within chromatin domains.

**Movie S7.**

MD simulation of 108-mer nucleosome (195 bp NRL) cluster plus 108 Linker Histone H1. As the simulation progresses, the nucleosome cluster adopts a more irregular structure, with H1 dynamically diffusing within the cluster (see Figs. 5A and S6A).

## Notes

### Competing Interest Statement

The authors have declared no competing interest.

### Summary of Updates

Figure 1 and Figure 5 were revised to clarify the multivalent interactions of H1; a biallelically tagged H1.2 cell line was examined.

